# HIV-1 Vpr induces cell cycle arrest and enhances viral gene expression by depleting CCDC137

**DOI:** 10.1101/2019.12.24.888230

**Authors:** Fengwen Zhang, Paul D. Bieniasz

## Abstract

The HIV-1 Vpr accessory protein induces ubiquitin/proteasome-dependent degradation of many cellular proteins by recruiting them to a cullin4A-DDB1-DCAF1 complex. In so doing, Vpr enhances HIV-1 gene expression and induces (G2/M) cell cycle arrest. However, the identities of Vpr target proteins through which these biological effects are exerted are unknown. We show that a chromosome periphery protein, CCDC137/cPERP-B, is targeted for depletion by HIV-1 Vpr, in a cullin4A-DDB1-DCAF1 dependent manner. CCDC137 depletion caused G2/M cell-cycle arrest, while Vpr-resistant CCDC137 mutants conferred resistance to Vpr-induced G2/M arrest. CCDC137 depletion also recapitulated the ability of Vpr to enhance HIV-1 gene expression, particularly in macrophages. Our findings indicate that Vpr promotes cell-cycle arrest and HIV-1 gene expression through depletion of CCDC137.

---

Human and simian immunodeficiency viruses (HIV-1, HIV-2 and SIVs) encode several accessory proteins, among which the function of ∼14-kDa Viral Protein R (Vpr) remains enigmatic. Replication deficits of inconsistent magnitude are evident HIV-1 strains lacking Vpr, particularly in primary macrophages (*1–3*), while deletion of Vpr from SIVmac modestly attenuates pathogenesis (*4, 5*).

Both Vpr and the related HIV-2/SIV accessory protein, Vpx, bind to VprBP (DCAF1) and thereby recruit the cullin 4A-containing E3 ubiquitin ligase complex (CRL4) (*6–13*). CRL4 recruitment by virion-associated Vpx and some SIV Vpr proteins can induce the degradation of the antiviral protein SAMHD1 following viral entry (*14–16*), but HIV-1, HIV-2, and many SIV Vpr proteins do not exhibit this activity. Rather, HIV-1 Vpr mediated CRL4 recruitment has different biological effects including G2/M cell-cycle arrest (*2, 8-13, 17-19*) and activation of the ATR (ataxia-telangiectasia and Rad3-related)-mediated DNA damage response (DDR) (*20–22*). HIV-1 gene expression is also enhanced by Vpr in some contexts, and similarly enhanced in cells arrested in G2/M (*2, 23–25*). However, none of the previously identified Vpr target proteins (*26–29*) have been demonstrated to be responsible for these biological effects.

We searched for HIV-1 Vpr target proteins using a proximity-dependent method (*30*) that employed a biotin-ligase, BirA(R118G) fused an HIV-1_NL4-3_ Vpr bait. Multiple nuclear proteins were biotinylated in BirA(R118G)-Vpr, but not BirA(R118G) expressing, proteosome inhibitor-treated, cells (Fig. S1A, Data S1). Ki-67, a proliferating cell marker, was the top ‘hit’, with >90 Ki-67 peptides detected in replicate experiments. However, Vpr did not induce Ki-67 depletion (Fig. S1B, C) and Ki-67 depletion did not induce G2/M arrest (Fig. S1D, E), suggesting that it is not a Vpr target protein. Ki-67 recruits a group of chromosome periphery proteins (cPERPs), that localize within the nucleus, primarily the nucleolus, during interphase but are relocalized to chromosome peripheries during mitosis (*31, 32*). Therefore, we conducted a focused screen of candidate target proteins that were either prominent hits in the BirA(R118G)-Vpr screen (Data S1), were nucleolar or nuclear proteins, members of the cPERP group, and/or were reported to bind Ki-67. Of numerous candidates tested, Vpr only induced the depletion of cPERP-B, also termed CCDC137(*32*) (Fig. S2A-C). Co-expression of a barely detectable amount of Vpr resulted in the removal of larger quantities of CCDC137, underscoring the potency of Vpr-induced CCDC137 depletion (Fig. 1A,B, Fig. S2D). The proteasome inhibitor MG132 blocked Vpr-induced CCDC137 degradation (Fig. 1C) as did depletion of DCAF1 by RNA interference (Fig. 1D). A truncated N-terminal (residues 1-154), but not a C-terminal (residues 155-289), CCDC137 fragment was depleted by Vpr (Fig. S3A). Scanning mutagenesis revealed that CCDC137 residues 61 to 75 were important for Vpr-induced depletion (Fig. 1E, Fig. S3B) and alanine substitutions of an LxxLL motif (positions 228-232) through which CCDC137 binds nuclear receptors (*33*), also reduced Vpr-induced CCDC137 depletion (L/A, mutation, Fig. 1E). Combined substitution of CCDC137 residues 61-65 or 66-70, coupled with 228-232 caused CCDC137 to be substantially Vpr-resistant (Fig. 1E). As previously reported (*32*), CCDC137 localized to nucleoli during interphase (Fig. S4A) and this property was unaffected by the aforementioned Vpr resistance-inducing substitutions (Fig. S4B).

**Fig. 1.**
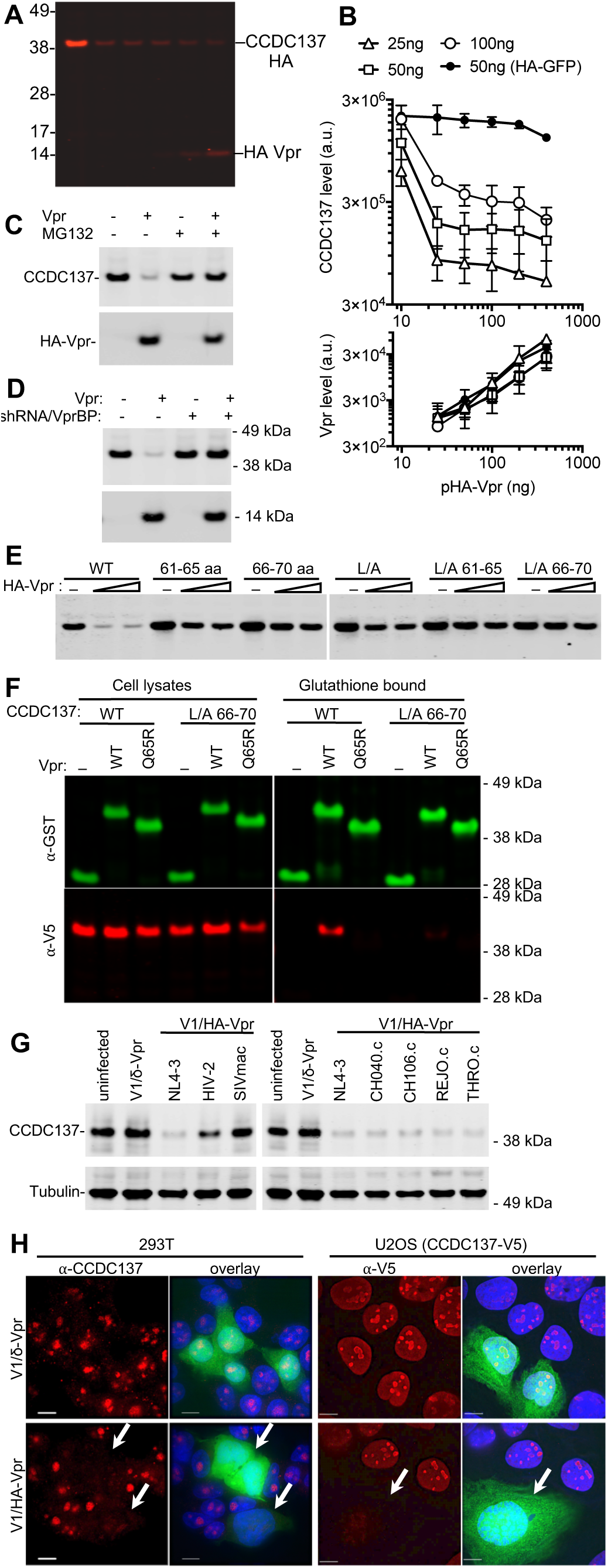
Binding to and depletion of CCDC137 by HIV-1 Vpr. (A) Western blot analysis of 293T cell lysates 28 hours after transfection of (100 ng ng/well) of an HA-CCDC137 expression plasmid and (0 ng, 25 ng, 50 ng, 100 ng, or 200 ng/well) of an HA tagged Vpr expression plasmid. (B) Western blot quantitation of HA-CCDC137 or HA-GFP, (upper panel) and HA-Vpr lower panel after transfection of 293T cells with the indicated amounts HA-Vpr (X-axis) or target protein expression plasmids. (C) Western blot analysis of 293T cell lysates at 28 hours after transfection with 150 ng of V5-tagged CCDC137 expression plasmid and 0 ng or 30 ng of HIV-1 Vpr expression plasmid. Cells were treated with 10 µM MG132 (+) or carrier (-) for 4 hours before harvest. (D) Western blot analysis of cell lysates from empty pLKO vector (-) or VprBP shRNA (+) transduced 293T cells after transfection of 150 ng of V5-tagged CCDC137 and 30 ng of HA-Vpr expression plasmid. (E) Western blot analysis of 293T cell lysates 28 hours after transfection with plasmids expressing HA-tagged CCDC137 encoding alanine substitutions at positions 61-65 or 66-70, in the LXXLL motif (L/A), or both, along with 0 ng, 50 ng or 100 ng of an HA-Vpr expression plasmid. (F) Western analysis of cell lysates and glutathione-agarose-bound fractions following transfection of 293T cells with plasmids expressing GST (-), GST-Vpr WT, or GST-Vpr mutant Q65R along with plasmids expressing HA-tagged wild-type (WT) or Vpr-resistant CCDC137 (L/A 66-70). (G) Western analysis of 293T cells 48 hours after infection (MOI = 2) with minimal HIV-1 viruses (V1) carrying no Vpr (V1/δVpr), or Vpr proteins from HIV-1_NL4-3_, HIV-2, SIVmac, or HIV-1 strains (CH040.c, CH106.c, REJO.c, or THRO.c). (H) Immunofluorescent detection of endogenous (293T cells, left) or V5-tagged (U2OS cells, right) CCDC137 at 48h after infection with V1/δ-Vpr (upper) or V1/HA-Vpr (lower). Infected, GFP-positive are indicated by arrows. Scale bar = 10µm.

Wild-type CCDC137, but not a Vpr-resistant CCDC137 mutant (L/A 66-70) could be co-precipitated with GST-Vpr (Fig. 1F) when coexpressed in 293T cells. Conversely, a mutant GST-Vpr (Q65R) that did not cause CCDC137 depletion (Fig. S5A) did not co-precipitate CCDC137 (Fig. 1F). Similarly, V5-tagged CCDC137 was co-immunoprecipitated with HA-tagged Vpr, unlike control nucleolus-associated V5-tagged control proteins (MKI67IP and RRP1, Fig. S5B). When the biotinylation proximity assay (*30*) was coupled to Western blot analysis, CCDC137 was found to be biotinylated by Vpr-BirA(R118G) but not BirA(R118G) (Fig. S5C).

We infected 293T or U2OS cells with a minimal version of HIV-1 (termed V1) in which only Tat, Rev, HA-tagged Vpr and GFP are expressed (Fig. S6). Infection with V1 lacking Vpr (V1/δ-Vpr) had no effect on endogenous CCDC137 levels, while infection with V1/HA-Vpr, encoding Vpr from one of several HIV-1 strains, caused CCDC137 depletion (Fig. 1G). HIV-2 Vpr induced partial CCDC137 depletion while SIV_MAC_ Vpr did not affect CCDC137 levels (Fig. 1G). Immunofluorescent staining of endogenous CCDC137 in 293T cells, or ectopically expressed V5-tagged CCDC137 in U2OS cells, showed that CCDC137 was depleted in V1/HA-Vpr infected (GFP+) cells compared to neighboring, uninfected (GFP-) cells or V1/δ-Vpr infected cells (Fig. 1H, Fig. S7A, B).

Lentiviral constructs expressing CAS9 and CCDC137-targeting CRISPR guide RNAs efficiently generated CCDC137 knockout alleles, but most transduced cells died and none of the numerous surviving cell clones that were analyzed contained frameshifting indels in both copies of CCDC137, suggesting that CCDC137 is essential for proliferating cell viability. Two CCDC137-targeting lentiviral shRNA vectors caused effective short term CCDC137 depletion after puromycin selection of 293T cells, with CCDC137/shRNAII causing more profound depletion than CCDC137/shRNAI (Fig. 2A). Notably, CCDC137/shRNA transduced cells accumulated in G2/M (Fig. 2B) and the extent of CCDC137 depletion and G2/M accumulation were correlated (Fig. 2A, B). A CCDC137 cDNA construct lacking the 3’UTR targeted by CCDC137/shRNAII substantially rescued G2/M arrest (Fig. 2B). G2/M accumulation was also evident upon CCDC137 depletion in several U2OS cell clones expressing an mClover or mKusabira-Orange2 (mKO2) fluorescent proteins fused to geminin (1-110 aa) that are depleted during G1 but present during G2/M (Fig. 2C, Fig S8). Accordingly, live cell imaging of U2OS/mClover-hGeminin (1-110 aa) cells revealed fluctuating fluorescence that disappeared upon cell division, while CCDC137 depleted cells did not divide and retained fluorescence, indicating G2/M growth arrest until apparent cell death (Movie S1). CCDC137 depletion also caused accumulation of nuclear foci of Ser139-phosphorylated histone H2A variant H2AX (γ-H2AX) (Fig. 2D), mimicking the reported Vpr-induced DDR (*21*). The CCDC137 depletion-induced DDR was rescued by expression of a CCDC137 cDNA (Fig. 2D).

**Fig 2.**
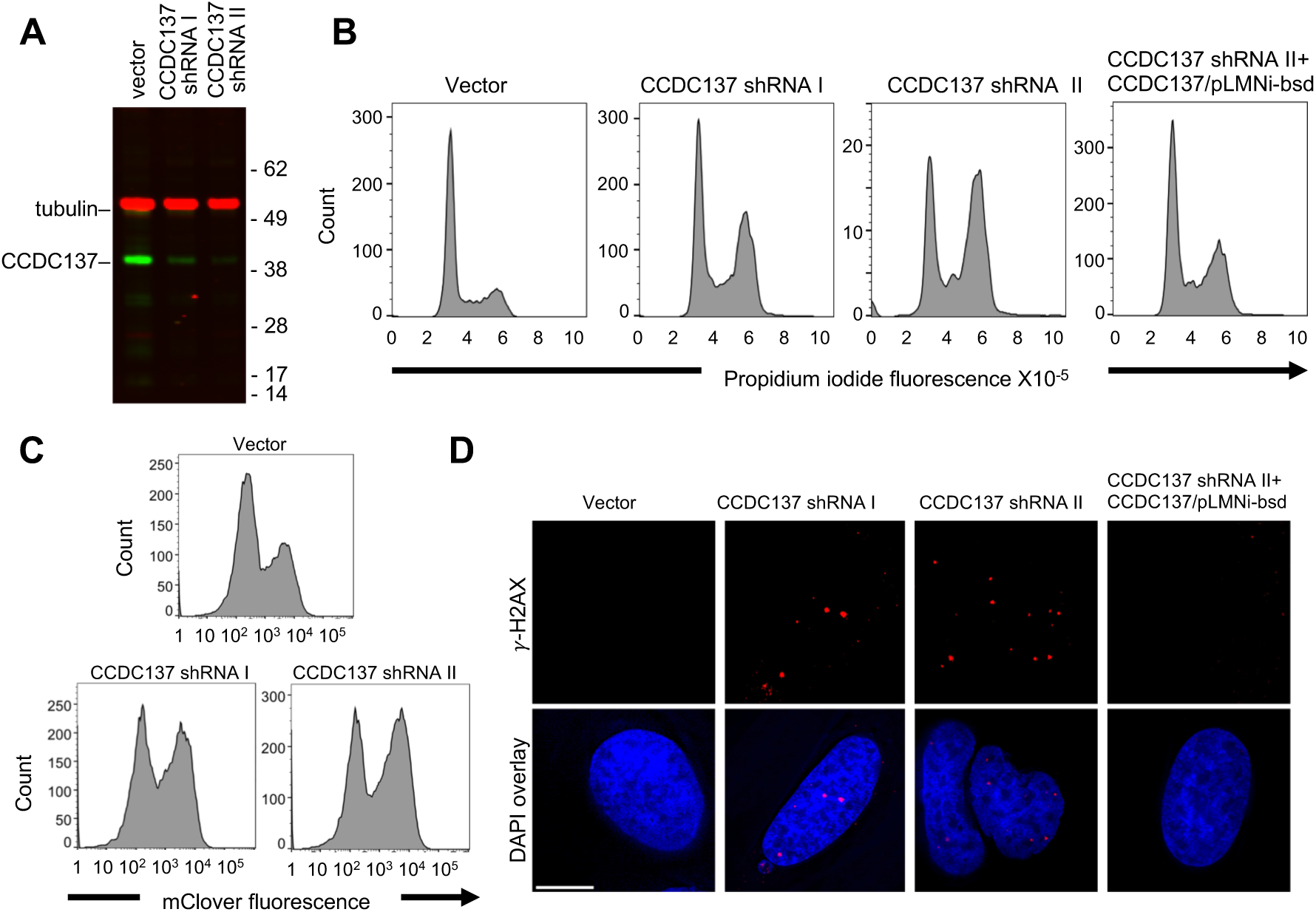
Induction of cell cycle arrest and DNA damage response by depletion of CCDC137. (A) Western blot analysis of 293T cells after transduction with pLKO lentivirus vectors carrying no shRNA (vector) or an shRNAs targeting one of two sequences on CCDC137 mRNA (I, II) and selection with puromycin for 40 hours. (B) Propidium iodide (PI) DNA content staining of 293T cells transduced with shRNA-carrying lentiviruses as in (A). Rightmost panel, 293T cells were also transduced with a retroviral vector expressing exogenous CCDC137. (C) FACS analysis of a U2OS-derived cell clone stably expressing mClover-hGeminin(1-110 aa), following transduction and selection as in (A). (D) Immunofluorescent staining of γ-H2AX foci (red) in U2OS cells following transduction as in (B). Scale bar: 10µm.

In U2OS cells containing doxyxcycline inducible CCDC137 constructs, wild-type CCDC137 was depleted following V1/HA-Vpr infection. Conversely, mutant CCDC137 (L/A 61-65 or L/A 66-70) largely resisted Vpr-induced depletion (Fig. 3A). Crucially, overexpression of WT CCDC137 partly ameliorated the G2/M arrest effect of Vpr, while the Vpr-resistant CCDC137 (L/A 66-70) mutant conferred nearly complete resistance to Vpr-induced G2/M arrest (Fig. 3B).

**Fig. 3.**
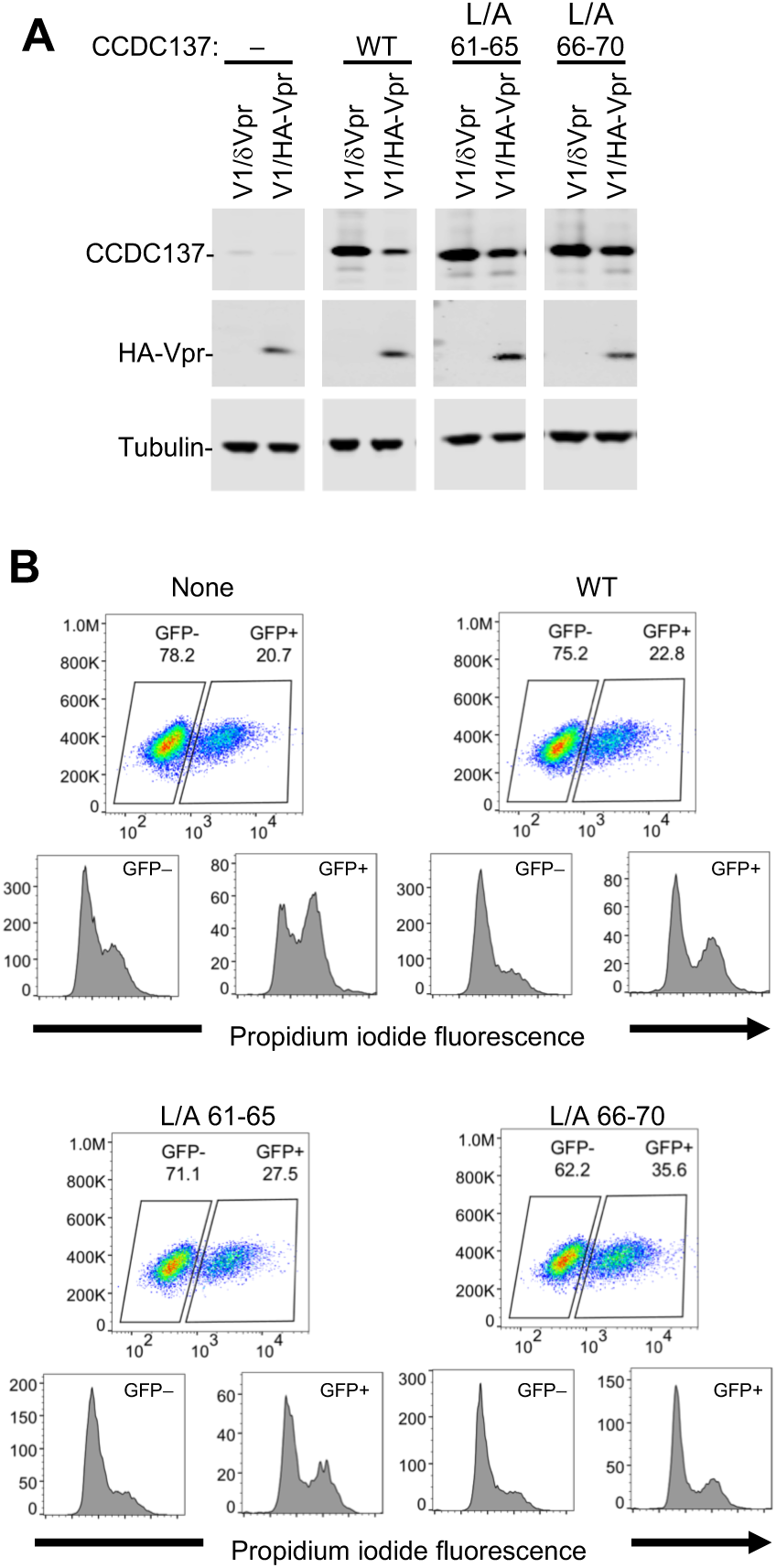
Vpr-resistant CCDC137 attenuates Vpr-induced G2/M arrest. (A) Western blot analysis of U2OS cells stably expressing doxycycline-inducible wild-type or Vpr depletion-resistant CCDC137 after doxycycline treatement for 24 hours and infection (MOI=2) with V1/HA-Vpr or V1/δ-Vpr for 48h. (B) DNA content assay of WT or mutant CCDC137-expressing cells infected (MOI=0.3) with V1/HA-Vpr. Upper plots depict forward scatter (Y axis) vs GFP fluorescence (X-axis) and percentages of GFP+ and GFP-cells. Lower plots depict PI staining in the uninfected (GFP-, lower left) and infected (GFP+, lower right) populations.

We assessed the effects of Vpr expression or CCDC137 depletion on HIV-1 gene expression in various cell types. V1/δ-Vpr/mCherry infected U2OS/mClover-hGeminin cells exhibited fluctuating mClover-hGeminin fluorescence indicative of a normal cell cycle, while V1/HA-Vpr/mCherry infection induced accumulation of mClover fluorescence in infected, mCherry+ cells (Movie S2). Notably, the level of mCherry fluorescence was clearly greater in the presence of Vpr (Movie S2). Moreover, FACS analysis or live-cell imaging showed that CCDC137 depletion using shRNA recapitulated the effect of Vpr, causing increased HIV-1 gene expression in V1/δ-Vpr infected U2OS cells (Fig 4A, B, Movie S3).

**Fig. 4.**
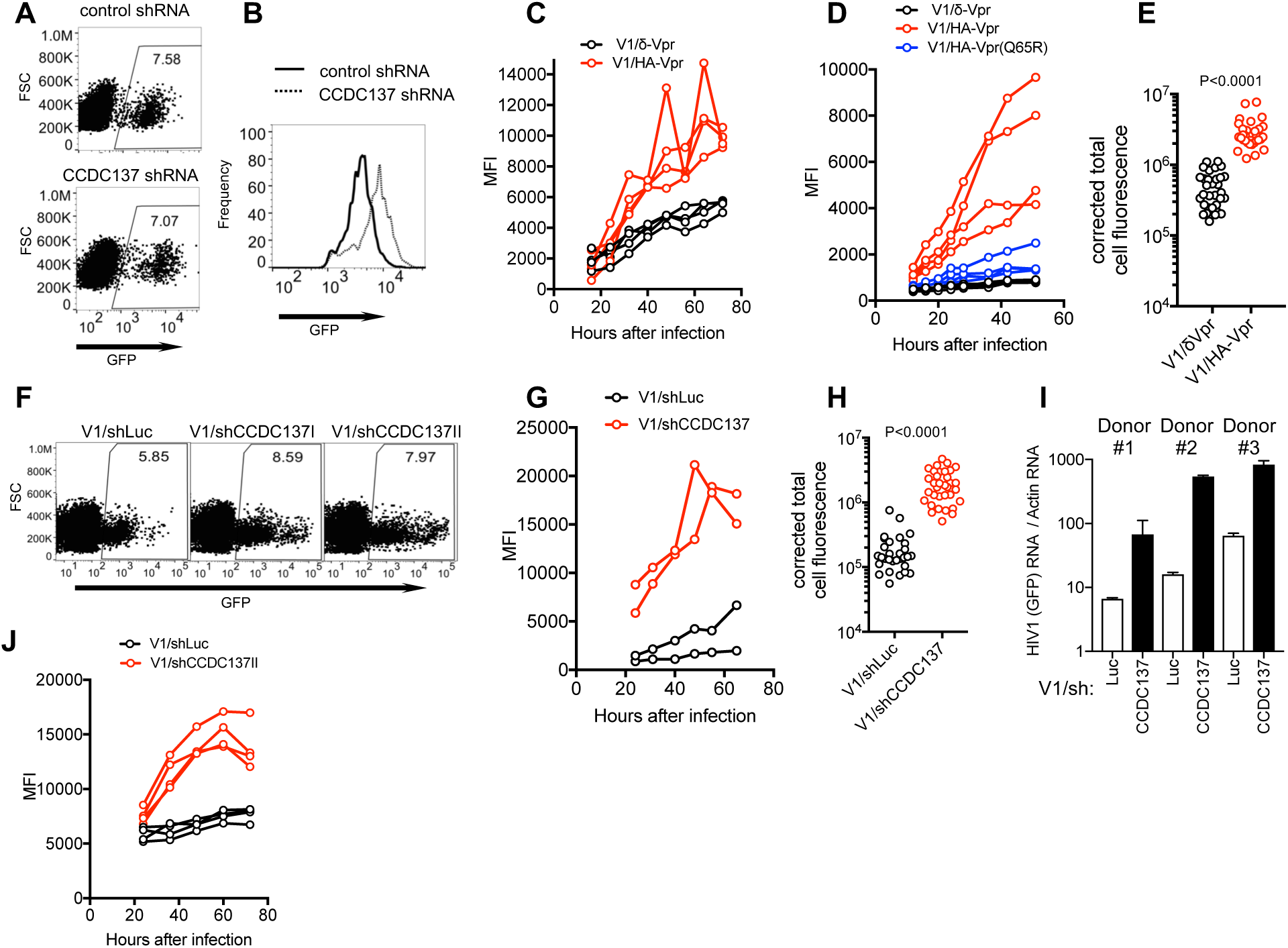
CCDC137 depletion increases HIV-1 gene expression. (A, B) HIV-1(GFP) expression in U2OS cells transduced with empty lentiviral vector (solid line) or CCDC137-targeting shRNA (dotted line) and selected with puromycin prior to infection with V1/δ-Vpr/GFP for two days. (B) represents histogram of GFP fluorescence in infected cells, gated as shown in (A), numbers in (A) represent % of cells within the gate. (C) GFP levels (mean fluorescence intensity (MFI), gated on infected cells) in primary CD4+ cells from four donors after infection with V1/δ-Vpr or V1/HA-Vpr. (D) GFP levels (MFI, gated on infected cells) in macrophages from four donors after infection with V1/δ-Vpr or V1/HA-Vpr. (E) GFP RNA levels (total cell fluorescence) determined by FISH analysis of macrophages infected with V1/δ-Vpr or V1/HA-Vpr. Each symbol is represents a single cell from a representative donor. (F, G) GFP expression in macrophages after infection with V1/shLuc or V1/shCCDC137. A representative donor (F), and MFI of infected cells for 2 donors (G), gated as in (F). Numbers in (F) represent % of cells within the gate. (H) GFP RNA levels determined as in (E) for macrophages infected with V1/shLuc or V1/shCCDC137. (I) RT-PCR measurement of HIV-1 RNA levels in 3 macrophage donors after infection with V1/shLuc or V1/shCCDC137II. (J) GFP levels (MFI, gated on infected cells) primary CD4+ cells from four donors after infection with V1/shLuc or V1/shCCDC137.

We examined effects of Vpr on gene expression in primary CD4+ T-cells and macrophages. Infection of primary CD4+ T-cells with V1/HA-Vpr resulted in modest (∼2-fold) higher levels of GFP expression than did infection with V1/δ-Vpr (Fig. 4C, Fig S9). Microscopic evaluation of V1 infected primary macrophages showed effective depletion of CCDC137 (Fig S10), and the effect of Vpr on HIV-1 gene expression was more pronounced therein, even though marked donor-to-donor variation was evident (Fig. 4F, Fig S10, S11A, Movie S4, S5). Several different Vpr proteins increased HIV-1 gene expression in macrophages (Fig. S11B) and similar Vpr-induced enhancement of HIV-1 gene expression was evident in macrophages infected with a full-length reporter virus (HIV-1_NHG_, Fig S11C, D), as previously reported (*2*). The Vpr-induced increase in GFP levels in V1 infected macrophages was accompanied by elevated HIV-1 RNA levels, as assessed by in-situ hybridization (Fig 4E, Fig S11E).

To enable Vpr-independent CCDC137 depletion in infected primary cells, while simultaneously measuring HIV-1 gene expression, we constructed a derivative of V1/δ-Vpr carrying an shRNA expression cassette (V1/sh, Fig S6). Infection with V1/sh enabled effective Vpr-independent CCDC137 depletion in infected macrophages (Fig S12). Strikingly, Vpr-independent CCDC137 depletion in V1/sh infected macrophages caused pronounced enhancement of HIV-1 reporter gene expression (Fig. 4F, G, Fig. S12, S13A, Movie S6, S7). The enhancing effect of CCDC137 depletion on HIV-1 gene expression was similarly evident when HIV-1 RNA levels were assessed by in-situ hybridization (Fig 4H, Fig S13B) or qRT-PCR (Fig. 4I) assays, indicating that CCDC137 depletion recapitulated the effect of Vpr on HIV-1 transcription or RNA stability. A more modest enhancement of HIV-1 gene expression in primary CD4+ T-cells was also evident upon CCDC137 depletion in the context of V1/sh (Fig 4J, Fig S13C). Overall, in three different cell types, shRNA-driven CCDC137 depletion had similar effects on HIV-1 gene expression as did Vpr.

Vpr-defective viruses are often selected in long-term HIV-1 passage or in chronically infected cells in vitro (*17, 18*). The seemingly deleterious effect of Vpr in these contexts is likely the result of G2/M arrest-induced reduction in cell longevity and, thus, viral burst size (*23*). Conversely, the life span of infected T-cells in vivo is likely limited by factors other than Vpr and even a modest Vpr-induced enhancement in viral gene expression should confer advantage. Macrophages are not cycling cells, and lack ATR, Rad17 and Chk1(*34*). Thus, G2/M arrest and DDR induction by Vpr are likely not relevant therein. The Vpr-induced enhancement of HIV-1 gene expression is particularly evident in macrophages (*2*). We surmise that the key consequence of Vpr-induced CCDC137 depletion is enhancement of HIV-1 gene expression, with G2/M arrest and DDR in cycling cells representing a side effect.

That Vpr associates with a cPERP protein is consistent with prior findings that Vpr binds to chromatin and, with VprBP, forms chromatin-associated nuclear foci (*35, 36*). At present, little is known about CCDC137 functions. CCDC137 can sequester the retinoic acid receptor (RAR) to the nucleolus (*33*) but whether this property is relevant to HIV-1 gene expression is unknown. Further work will be required to discern the mechanistic details of how CCDC137 affects G2/M transition and inhibits HIV-1 gene expression.

## Acknowledgments

We thank Proteomics Resource Center, Rockefeller University for mass spectrometry analysis. We thank Agata Smogorzewska, Theodora Hatziioannou and Trinity Zang for reagents and other members of the Bieniasz and Hatziioannou laboratories for helpful discussions.

## Author contributions

FZ and PB designed the research and wrote the paper. FZ conducted the experiments.

## Data and materials availability

All data are available in the manuscript or the supplementary materials. Requests for materials should be addressed to P.D.B.

## Materials and Methods

### Antibodies

Monoclonal antibodies used herein included anti-FLAG (Sigma), anti-HA (BioLegend anti-HA.11), anti-HIV capsid p24 (183-H12-5C, NIH AIDS Research and Reference Reagent Program), anti-Ki67 (abcam, ab16667), anti-tubulin (Sigma, T9026), and anti-V5 (Invitrogen, R960-25). Polyclonal antibodies were purchased from ThermoFisher (anti-V5, PA1-993), or Abcam (anti-CC137, ab185368 or ab183864; anti-gamma H2AX phosphor S139, ab11174; anti-Fibrillarin, ab5821). Secondary antibodies included goat anti-mouse or anti rabbit IgG conjugated to Alexa Fluor 488 or Alexa Fluor 568 (Invitrogen) for immunostaining, or IRDye® 800CW and IRDye® 680, or IRDye 680RD Streptavidin (LI-COR Biosciences) for Western blot analysis.

### Plasmid Construction

An HA-epitope was fused in-frame at the N-terminus of HIV-1_NL4-3_ Vpr and subcloned into pCR3.1 for transient cotransfection experiments. To express Vpr driven by an HIV-1 long terminal repeat (LTR) in the context of a viral genome, a proviral plasmid was generated harboring a minimal viral genome (V1) (*37*), engineered from HIV-1_NL4/3_ (R7/3) harboring large deletions or inactivating mutations in (Gag, Pol, Vif, Vpu and Env) and in which Nef is replaced with GFP. Sequences encoding an HA epitope were fused in frame at N-terminus of Vpr to generate V1/HA-Vpr while V1/δ-Vpr was constructed by deletion of the nucleotide sequences between the cPPT/FLAP and the SalI site within Vpr. In some constructs, the WT HIV-1_NL4-3_ Vpr was replaced with Vpr-encoding sequences from Q65R NL4-3 Vpr, transmitted HIV-1 founder strains, HIV-2, SIV_AGM_ Sab, or SIVmac. To construct the V1/HA-Vpr or V1/δ-Vpr expressing mCherry (V1/mCherry), an open reading frame encoding mCherry was amplified, digested with NotI/XhoI, and inserted into V1 to replace GFP. To construct the V1/shCCDC137II carrying an shRNA targeting CCDC137, the DNA sequence containing the U6 promoter and shRNA targeting CCDC137 (I: CAGATGCTGCGGATGCTTCT; II: GGTGAAACATGATGACAACA) was PCR amplified and, after digestion with KpnI, subcloned into V1 carrying inactivating mutations at the 5’ end of Vpr (ATGGAACAA/GTGGAATAA), upstream 5’ of the RRE (Fig. S6). The control V1/shLuc vector contained the DNA sequence carrying U6 promoter-shRNA targeting luciferase (CGCTGAGTACTTCGAAATGTC) at the same position. V1 derivatives expressing BirA (R118G) and BirA (R118G)-Vpr were constructed by insertion of nucleotide sequences encoding BirA (R118G) or BirA (R118G)-Vpr fusion proteins into the GFP (Nef) position in V1 vector.

Plasmids expressing the various human proteins (CREB1, CREB3L1, hnRNP D, hnRNP F, POC1A, PPM1G, CCDC137, PES1, WDR18, WDR74, MAK16, NOC2L, RRS1, NIP7, hnRNPA1, hnRNP H, hnRNP K, hnRNP R, hnRNP U, hnRNP C, hnRNP A2B1, and DHX9) with a V5 epitope fused to the C-terminus were from a pCSGW-based human ORF lentiviral library. Plasmid pCR3.1 was used to express Bop1, NCL, MKI67IP, or WDR12 with an Flag epitope fused to their N-termini or to express hnRNP A2B1 or DHX9 with an HA epitope fused to their N-termini. Subsequently, CCDC137 was subcloned into pLNCX2 expression vector with an HA epitope fused to its C-terminus. CCDC137 alanine scanning mutants were generated by overlap-extension PCR amplification, using the wild-type CCDC137-HA expression plasmid as the template and inserted into the pLNCX2 expression vector. All aforementioned plasmids were constructed using PCR, and sequences for cDNA and oligonucleotides are available upon request.

GST-Vpr expression plasmids were based on pCAGGS and were constructed by PCR amplification of the Vpr coding region from HIV-1_NL4-3_ which was then inserted in-frame at the 3’ end of GST encoding sequences.

To construct a tetracycline-inducible CCDC137 expression vector, wild-type and mutant CCDC137 were amplified, using the corresponding pCR3.1 expression plasmids as templates, and inserted into LKO-derived lentiviral expression vector pLKOΔ-puro (*38*) which also included a puromycin resistance cassette. All cloned coding sequences were verified by DNA sequencing, oligonucleotide sequences used in construction are available upon request.

For knockdown experiments, the lentiviral vector pLKO.1-TRC (*39*) was used to deliver shRNAs. For CCDC137 shRNAs, the lentiviral vector contains two functional elements, shRNA targeting sequences (I: CAGATGCTGCGGATGCTTCT; II: GGTGAAACATGATGACAACA) and puromycin-resistance cassette.

### Cell lines

Human embryonic kidney HEK-293T were maintained in DMEM supplemented with 10% fetal bovine serum (Sigma F8067) and gentamycin (Gibco). Human bone osteosarcoma epithelial cells (U2OS, ATCC® HTB-96™) were grown in McCoy’s 5a Medium Modified (ATCC® 30-2007™) /10%FCS/gentamycin. T lymphocyte cell lines CEM, Jurkat, MT4 and SupT1 were maintained in RPMI supplemented with 10% fetal bovine serum (FCS) and gentamycin. All cell lines used in this study were monitored by SYBR® Green real-time PCR RT assay periodically to ensure the absence of retroviral contamination. To construct cell-cycle reporter cell line, U2OS cells were transduced with a retroviral vector (pLHCX) encoding Clover (a rapidly-maturing green/yellow fluorescent protein) or mKusabira-Orange2 (mKO2, an orange fluorescent protein) fused to the N-terminus of Geminin 1-110 aa. Single cell clones were isolated after hygromycin selection.

### Primary cells

Human lymphocytes were prepared from Leukopaks from NY Blood Center by spinning on top of lymphocyte separation medium (Corning). Macrophages were then isolated by plastic adherence and differentiated using GM-CSF (Thermo Fisher). CD4+ T cells were isolated using an EasySep kit (StemCell) and maintained in RPMI/10%FCS supplemented with IL2.

### Transfection experiments

For transfection experiments in 293T cells, cells were seeded at a concentration of 1.5×10^5^ cells/well (24-well plate), 3×10^5^ cells/well (12-well plate) or 2×10^6^ (10-cm dish) and transfected on the following day using polyethylenimine (PolySciences).

To test whether Vpr could induce depletion of proteins, 293T cells in 24-well plates were transfected with 200 ng of pCR3.1-based plasmids expressing Flag-tagged proteins or HA-tagged proteins, or pCSGW-based plasmids expressing V5-tagged proteins (from human ORFs lentiviral library), along with increasing amounts (0 ng, 25 ng, or 50 ng) of a pCR3.1/HA-Vpr expression plasmid. The total amount of DNA was held constant by supplementing the transfection with empty expression vector. Cells were harvested at 28 hours post transfection and subjected to Western blot analysis.

To assess the potency with which HIV-1 Vpr induced CCDC137 depletion, 293T cells in 24-well plates were transfected with varying amounts (0 ng, 100 ng, 200 ng, or 400 ng/well) of a pCR3.1/CCDC137-HA expression plasmid and increasing amounts (0 ng, 25 ng, 50 ng, 100 ng, or 200 ng/well) of a pCR3.1/HA-Vpr expression plasmid. The total amount of DNA was held constant by supplementing the transfection with empty expression vector.

### Generation of HIV-1 and lentiviral vector stocks

To generate V1-derived viral stocks, 293T cells were transfected with 5 µg of pV1-derived proviral plasmids encoding no Vpr or the various HA-tagged Vpr proteins, 5 µg of an HIV-1 Gag-Pol expression plasmid (pCRV1/GagPol) and 1 µg of VSV-G expression plasmid into 293T cells in 10-cm dishes. Virus-containing supernatant was collected and filtered (0.2µm) 2 days later. The lentiviral vectors that transduced CCDC137 cDNAs or shRNAs were similarly generated, except that pLKOΔ-puro or pLKO.1-TRC-derived plasmids were used in place of pV1-derived plasmids.

### Identification of Vpr proximal and interacting proteins

Ten million MT4 cells were transduced with V1-based constructs expressing BirA (R118G) or BirA (R118G)-Vpr at an MOI of 2, treated with 50 µM biotin (Sigma) and, after treatment with 10 µM MG132 for 4 hours, cells were harvested 48 hours after transduction. Biotinylated proteins were purified using a previously described protocol (*30*). In brief, cells were lysed in lysis buffer (50 mM Tris, pH 7.4, 500 mM NaCl, 0.4% SDS, 5 mM EDTA, 1 mM DTT, and 1x complete protease inhibitor [Roche]) and sonicated. After addition of Triton X-100 and further sonication, cell lysates were centrifuged and cleared supernatants were incubated with Dynabeads (MyOne Steptavadin C1 [Invitrogen]) for 4 hours. Beads were then washed with 2% SDS, followed by wash thoroughly with buffer 2 (0.1% deoxycholate, 1% Triton X-100, 500 mM NaCl, 1 mM EDTA, and 50 mM Hepes, pH 7.5), buffer 3 (250 mM LiCl, 0.5% NP-40, 0.5% deoxycholate, 1 mM EDTA, and 10 mM Tris, pH 8.1) and buffer 4 (50 mM Tris, pH 7.4, and 50 mM NaCl). Biotinylated proteins were eluted from the beads with NuPAGE® LDS sample buffer supplemented with 200 µM biotin and separated by NuPAGE Bis-Tris Gels. Each gel lane was cut in 5 bands and subjected to LC-MS/MS analysis (Proteomics Resource Center, Rockefeller University).

### Immunoprecipitation

HEK-293T cells were transiently transfected with plasmids expressing HA-Vpr and V5-tagged human protein factors, and treated with 10 µM MG132 for 4 hours before harvest and lysis with ice-cold lysis buffer (50 mM Tris, pH 7.4, 150 mM NaCl, 0.5 mM EDTA, 1% digitonin [Sigma], supplemented with 1X complete protease inhibitor [Roche]). After lysis on ice for 10 min, followed by centrifugation at 10,000 rpm for 10 min at 4°C, clarified lysates were mixed with 1 µg anti-HA monoclonal antibody and rotated with 30 µl pre-equilibrated Protein G Sepharose 4 Fast Flow resin (GE healthcare) for 3 hours at 4°C. The resin was then washed 3 times with wash buffer (50 mM Tris, pH 7.4, 150 mM NaCl) and the bound proteins were eluted with SDS-PAGE sample buffer and analyzed by Western blotting.

### Glutathione-S-Transferase (GST) fusion protein interaction assay

Human 293T cells in 6-well plates were co-transfected with 100 ng of GST or 1 µg of GST-Vpr expression plasmids and 500 ng of HA-tagged CCDC137 expression plasmids. The total amount of DNA was held constant by supplementing the transfection with empty expression vector. Two days later, cells were treated with MG 132 for 4 hours and then lysed in Lysis buffer (50 mM Tris, pH 7.4, 150 mM NaCl, 5 mM EDTA, 5% glycerol, 1% Triton X-100, and 1x complete protease inhibitor [Roche]). Cleared lysates were then incubated with glutathione sepharose (GE healthcare) for 4 hours at 4°C and, after wash with buffer (50 mM Tris, pH 7.4, 150 mM NaCl, 5 mM EDTA, 0.1% Triton^TM^ X-100), bound proteins were eluted in SDS-PAGE sample buffer and subjected to Western blot analysis.

### BirA-fusion protein interaction assay

Human 293T cells in 6-well plates were transfected with a V5-tagged CCDC137 expression plasmid and BirA (R118G) or BirA (R118G)-Vpr expression plasmid. Cells were treated with 50 µM biotin at 24 hours after transfection, and 10 µM MG132 at 40 hours post transfection, and harvested at 44 hours post transfection. Cells were then lysed and cleared lysates were incubated with Dynabeads (MyOne Steptavadin C1) above. After thorough wash, biotinylated proteins were eluted from the beads with SDS-PAGE sample buffer supplemented with 200 µM biotin and subjected to Western blot analysis.

### Western blot analysis

Cell lysates and immunoprecipitates were separated on NuPage Novex 4-12% Bis-Tris Mini Gels (Invitrogen), and NuPAGE™ MES SDS running buffer (Invitrogen, NP0002) was used when Vpr was detected. Proteins were blotted onto nitrocellulose membranes. Thereafter, the blots were probed with primary antibodies and followed by secondary antibodies conjugated to IRDye 800CW or IRDye 680. Fluorescent signals were detected and quantitated using an Odyssey scanner (LI-COR Biosciences).

### Cell cycle analysis

To determine effects of Vpr on cell cycle, V1 based viral stocks were used to inoculate 2.5×10^5^ U2OS cells in 6-well plates at an MOI of 0.5 to 1. At 48 hours post-infection, cells were trypsinized, fixed with paraformaldehyde (PFA) in phosphate-buffered saline (PBS), washed with PBS, and fixed again in 70% ethanol. After an additional wash with PBS, the cells were resuspended in FxCycle^TM^PI/RNase Staining Solution (Invitrogen) and incubated at 30°C for 30 min. Flow cytometric analysis was performed using Attune® NxT Acoustic Focusing Cytometer (ThermoFisher Scientific).

Alternatively, cells were transduced with pLKO.1-TRC-derived vectors encoding shRNA targeting CCDC137. After 48 hours, cells were selected in 1 µg ml^−1^ puromycin or 5 µg ml^−1^ blasticidin prior to propidium iodide staining, 40 hours later as described above.

In some experiments, U2OS cells expressing mKO2-hGeminin (1-110 aa) or mClover hGeminin (1-110 aa) were transduced with LKO-derived lentiviral vectors encoding shRNAs targeting CCDC137. After 48 hours, cells were selected in 1 µg ml^−1^ puromycin prior to FACS analysis, 48 hours later.

For the experiment in Fig. 3, U2OS cells expressing doxycycline-inducible CCDC137 were generated by transduction with a LKO-derived lentiviral vector (*38*) followed by selection in 1 µg ml^−1^ puromycin. Cells were plated at the density of 2.5×10^5^ in 6-well plates in the presence of doxycycline and the next day were infected with V1/HA-Vpr at an MOI of 2 or 0.5. At 12 hours post-infection, doxycycline was replenished and at 48 hours post-infection, cells were harvested for Western blot analysis (for cells infected at high MOI) or cell cycle analysis (for cells infected at low MOI).

### Fixed cell microscopy

U2OS cells expressing V5-tagged CCDC137 were seeded on 3.5-cm, glass-bottomed dishes coated with poly-L-Lysine (MatTek). At 48 hours after infection with V1/δ-Vpr or V1/HA-Vpr, cells were then fixed with 4% paraformaldehyde, permeabilized with 0.1% Triton X-100 and incubated with mouse anti-V5 monoclonal antibody (Invitrogen) and rabbit anti-Fibrillarin polyclonal antibody (abcam) followed by goat anti-mouse IgG Alexa fluor-594 conjugate and goat anti-rabbit IgG Alexa fluor-488 conjugate (Invitrogen). Images were captured using an DeltaVision OMX SR imaging system (GE Healthcare). Endogenous CCDC137 in 293T cells or in macrophages was stained with rabbit anti-CCDC137 polyclonal antibody (abcam) using the same procedure.

For detection of γ-H2AX foci, U2OS cells were transduced with lentiviruses encoding CCDC137-targeting shRNAs and, after selection with puromycin, cells were seeded on 3.5-cm, glass-bottomed dishes coated with poly-L-Lysine (MatTek). Nuclear foci were visualized by immunostaining with rabbit anti-γ-H2AX (abcam) followed by a goat anti-rabbit IgG Alexa Fluor-594 conjugate (Invitrogen). A Z-series of images were acquired using an DeltaVision OMX SR imaging system (GE Healthcare).

### Fluorescence in situ hybridization (smFISH)

At 48 h postinfection, macrophages in 8-well chamber slides were washed with PBS (Ambion), fixed with 4% formaldehyde (Thermo Fisher) in PBS and permeabilized with 70% ethanol. Then, the cells were washed with Stellaris RNA FISH wash buffer A (Biosearch Technologies) and GFP RNA detected by incubation using 0.125 µM Cy5-labeled probes in Stellaris RNA FISH hybridization buffer (Biosearch Technologies) for 18 h at 37°C. The cells were then washed twice in Stellaris RNA FISH wash buffer A (Biosearch Technologies) with the presence of Hoechst stain during the second wash. Then the cells were washed briefly with Stellaris RNA FISH wash buffer B (Biosearch Technologies), rinsed three times with PBS and subject to imaging by deconvolution microscopy (DeltaVision OMX SR imaging system). All images were generated by maximum intensity projection using the Z project function in ImageJ 1.52b. Corrected total cell fluorescence (CTCF) was calculated as the following: CTCF=Integrated Density - (Area of selected cell x Mean fluorescence of background readings). The 28 probes against GFP (cggtgaacagctcctcgc, gaccaggatgggcaccac, gtttacgtcgccgtccag, acacgctgaacttgtggc, gccggtggtgcagatgaa, ggtggtcacgagggtggg, actgcacgccgtaggtca, tcggggtagcggctgaag, agtcgtgctgcttcatgt, ggcatggcggacttgaag, ctcctggacgtagccttc, gccgtcgtccttgaagaa, tcggcgcgggtcttgtag, tgtcgccctcgaacttca, ctcgatgcggttcaccag, tgaagtcgatgcccttca, caggatgttgccgtcctc, cgttgtggctgttgtagt, gcttgtcggccatgatat, gtcctcgatgttgtggcg, gtagtggtcggcgagctg, cgatgggggtgttctgct, ttgtcgggcagcagcacg, gactgggtgctcaggtag, ttggggtctttgctcagg, catgtgatcgcgcttctc, ggtcacgaactccagcag, cttgtacagctcgtccat) were designed using the Stellaris Probe Designer, version 2.0 (Biosearch Technologies).

### Measurement of mRNA levels using qPCR

Macrophages (6 × 10^5^) were infected with V1/sh carrying shRNA targeting luciferase or CCDC137 (II), and 48 h later RNA was isolated using the NucleoSpin RNA Kit (Macherey-Nagel). RNA levels were determined with Power SYBR® Green RNA-to-CT™ 1-Step Kit using a StepOne Plus Real-Time PCR system (Applied Biosystems). For relative quantification, samples were normalized to actin. The sequences of primers are GAGCGCACCATCTTCTTCAA (GFP forward), TCCTTGAAGTCGATGCCCTT (GFP reverse), CATGTACGTTGCTATCCAGGC (actin forward) and CTCCTTAATGTCACGCACGAT (actin reverse). Relative GFP expression was calculated as the value of 2^-[ΔCt (GFP)- ΔCt (actin)].

### Live cell microscopy

To monitor cell cycle and HIV-1 (V1) gene expression infection in living cells, U2OS cells expressing mClover-hGeminin (1-110 aa) or primary macrophages were transferred into glass-bottom dishes and time-lapse microscopy was performed using a VivaView FL incubator microscope (Olympus). In some experiments, cells were transduced with lentiviruses containing shRNA targeting CCDC137, 36 hours prior to imaging. In some experiments, cells were infected with V1/δ-Vpr or V1/HA-Vpr expressing mCherry or GFP 12 to 24 hours prior to imaging.

Images were captured every 30 minutes using GFP, mRFP and DIC filter sets for up to 72 hours. Preparation of movies was done using MetaMorph software (Molecular Devices) as previously described (*40*). Images had a depth of 12 bits, i.e., an intensity range of 0–4095.

---

**Fig. S1.**
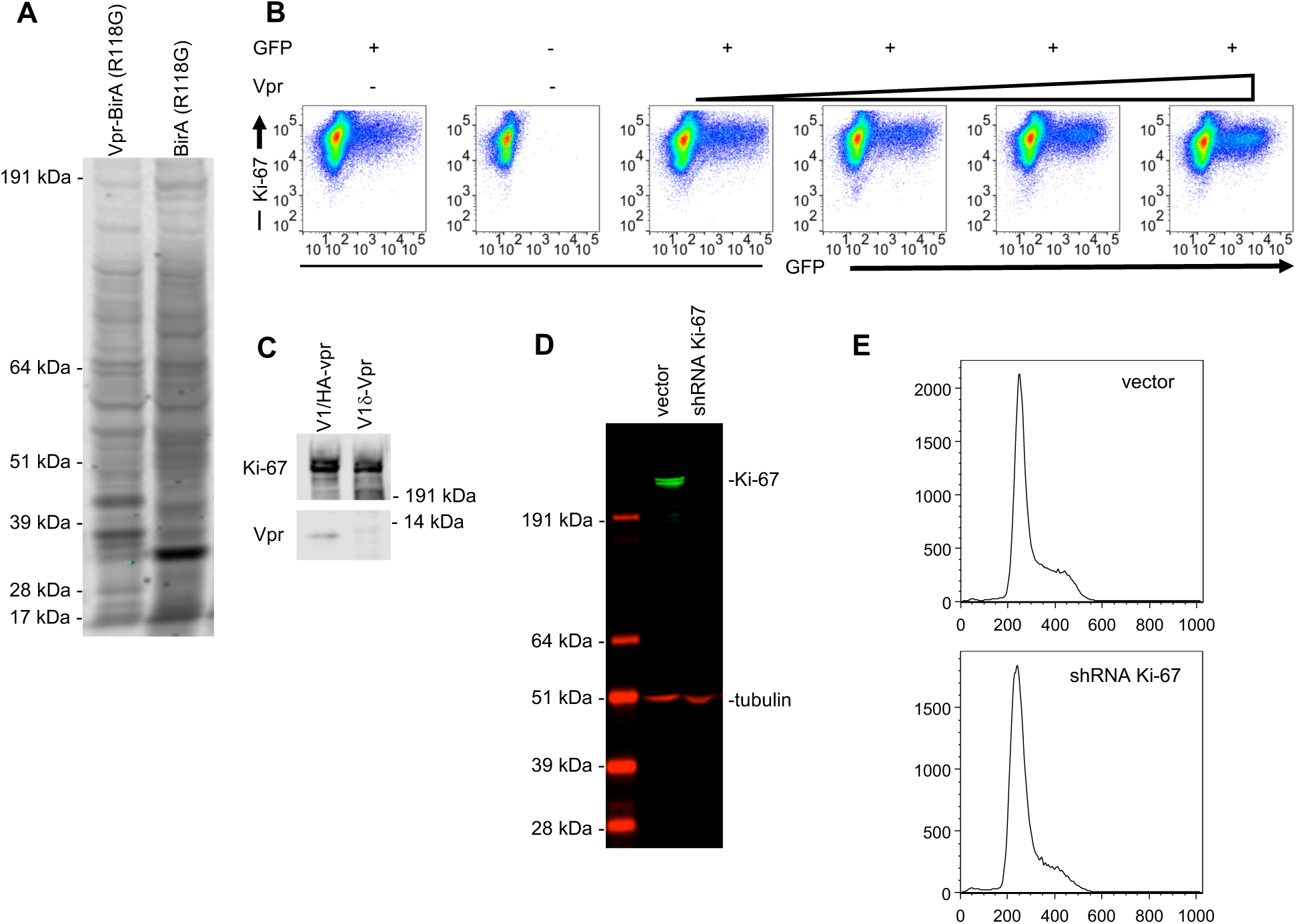
Discovery and analysis of proteins labelled with Vpr-BirA (R118G). (A) Proteins recovered from lysates of MT-4 cells expressing Vpr-BirA (R118G) or BirA (R118G) using streptavidin beads. Biotinylated proteins were detected with IRDye 680RD-conjugated Streptavidin. (B) Immunostaining/FACS analysis of endogenous Ki-67 levels in 293T cells 28 hours after transfection (in 6-well plates) with 100 ng of a GFP expression plasmid and increasing amounts (0 ng, 50 ng, 100 ng, 200 ng, or 400 ng) of an HA-Vpr expression plasmid. (C) Western blot analysis of endogenous Ki-67 and HA-Vpr expression in 293T cells 48 hours after infection with V1/δ-Vpr or V1/HA-Vpr at an MOI of 2. (D) Western blot analysis of Ki-67 and tubulin (loading control) levels in U2OS cells transduced with lentiviruses carrying no shRNA (vector) or shRNA against Ki-67 (shRNA Ki-67) after selection with puromycin. (E) Analysis of DNA content by propidium iodide staining in U2OS cells transduced with lentiviruses carrying no shRNA (vector) or shRNA against Ki-67 (shRNA Ki-67) after selection with puromycin.

**Fig. S2.**
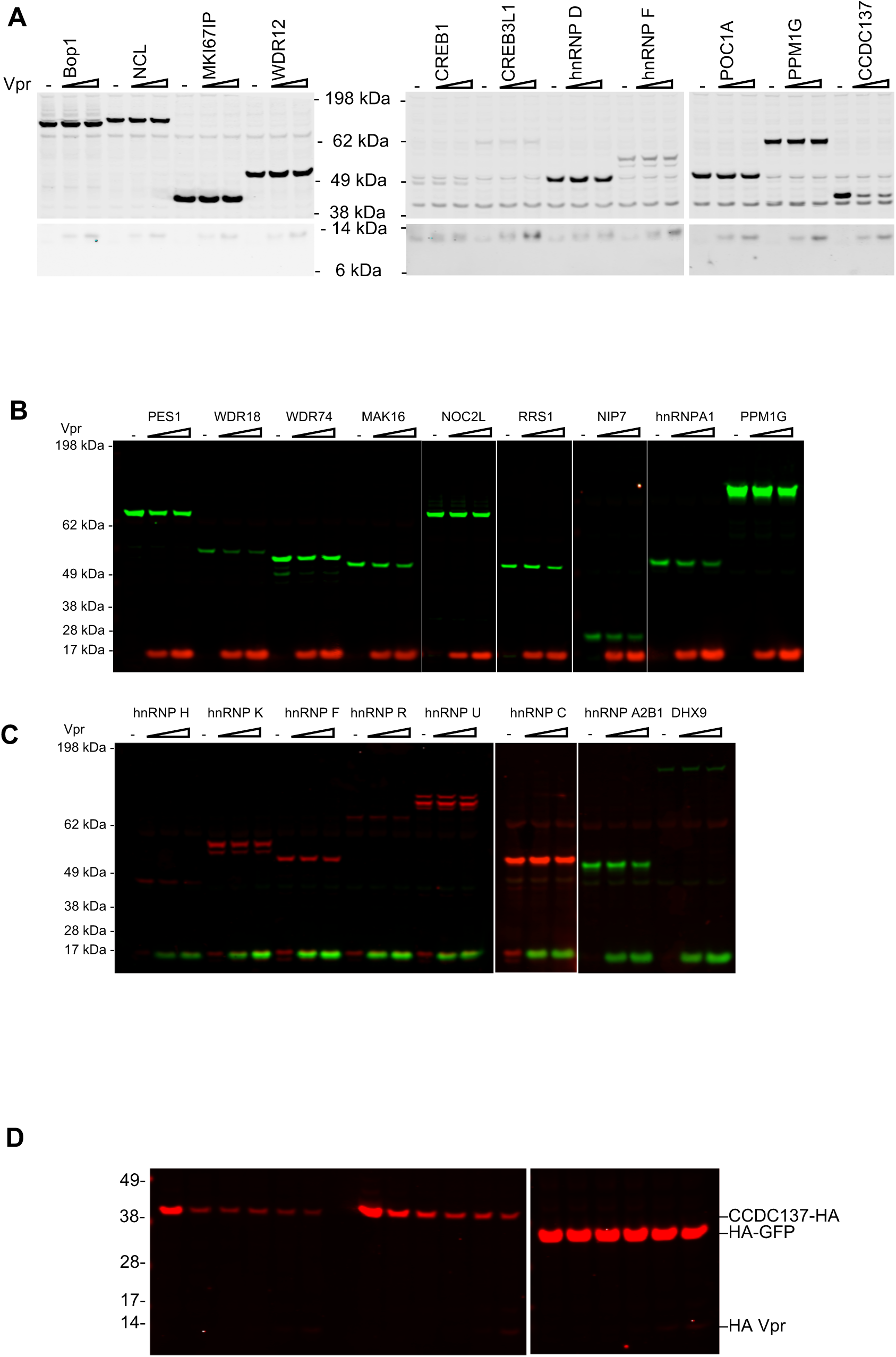
Focused screens for Vpr target proteins. (A) Western blot analysis of 293T cell lysates at 28 hours after transfection with 200 ng of plasmids expressing Flag-tagged Bop1, NCL, MKI67IP, or WDR12 (left panel), or V5-tagged CREB1, CREB3L1, hnRNPD, hnRNP F, POC1A, PPM1G, or CCDC137 (right panel) along with increasing amounts (0 ng, 25 ng, or 50 ng) of an HA-Vpr expression plasmid. (B) Western blot analysis of 293T cell lysates at 28 hours after transfection of 200 ng of plasmids expressing V5-tagged PES1, WDR18, WDR74, MAK16, NOC2L, RRS1, NIP7, hnRNPA1, or PPM1G (shown in green) along with increasing amounts (0, 100 ng, or 300 ng) of HA-Vpr expression plasmid (shown in red). (C) Western blot analysis of 293T cell lysates after transfection of plasmids expressing V5-tagged hnRNP F, hnRNP H, hnRNP K, hnRNP R, hnRNP U, or hnRNP C (shown in red) or HA-tagged hnRNP A2B1 or DHX9 (shown in green) with increasing amounts of HA-Vpr expression plasmid (shown in green). (D) Western blot analysis of 293T cell lysates 28 hours after transfection of varying amounts (0 ng, 100 ng, 200 ng, or 400 ng/well) of an HA tagged CCDC137 expression plasmid or 50 ng of HA tagged GFP expression plasmid and increasing amounts (0 ng, 25 ng, 50 ng, 100 ng, or 200 ng/well) of an HA tagged Vpr expression plasmid.

**Fig. S3.**
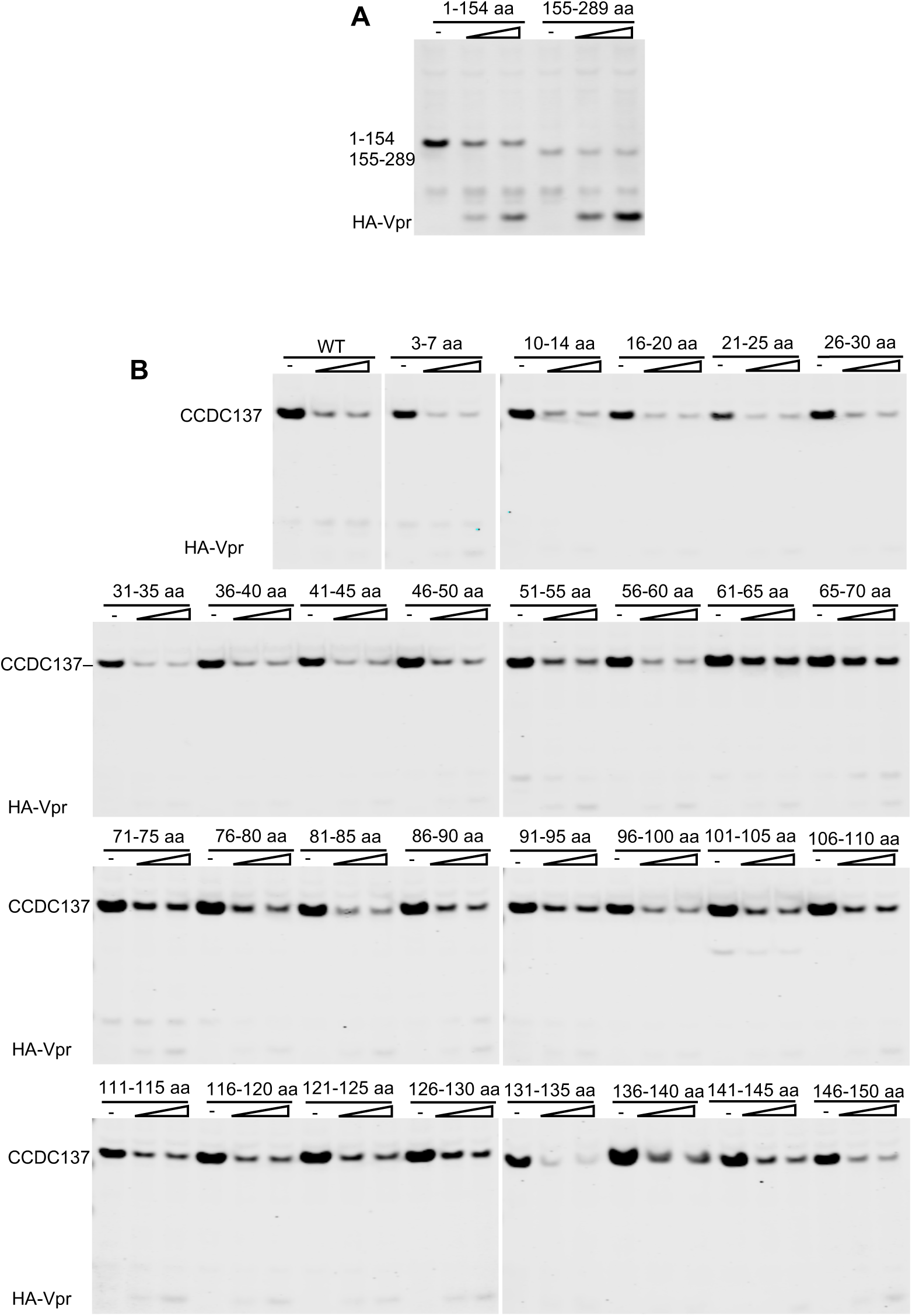
Mapping of degrons in CCDC137 for Vpr-induced depletion. (A) Western blot analysis (at 28 hours post transfection) following co-transfection of plasmids expressing truncated CCDC137-HA (namely 1-154 aa and 155-289 aa) with increasing amounts (0 ng, 50 ng, or 100 ng) of an HA-Vpr expression plasmid. (B) Western blot analysis (at 28 hours post transfection) following co-transfection of CCDC137-HA expression plasmids with increasing amounts (0 ng, 25 ng, or 50 ng) of HA-Vpr expression plasmids. The CCDC137 proteins encoded alanine scanning mutations at the indicated positions. Each indicated 5-aa stretch was replaced with Ala except where Ala was present in the original sequence.

**Fig. S4.**
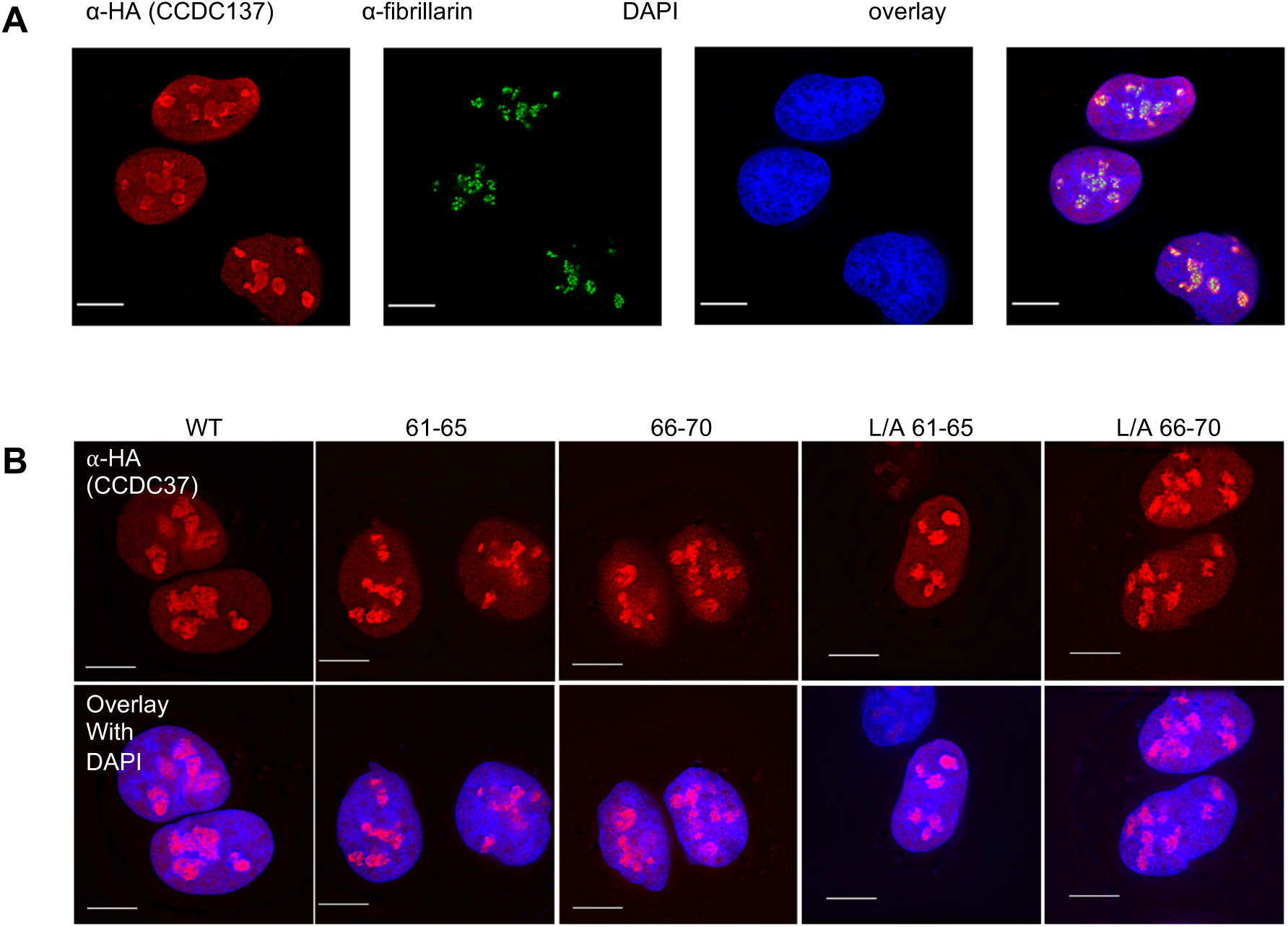
CCDC137 localization in the nucleolus. (A) Immunofluorescent staining of U2OS cells stably expressing V5-tagged wild-type CCDC137 (red). Endogenous fibrillarin was also immunostained (green) as was DNA (blue). Scale bar: 10µm. (B) Immunostaining of U2OS cells stably expressing HA-tagged wild-type (WT) or CCDC137 bearing Ala substitutions at positions 61 to 65 (61-65), 66 to 70 (66-70) alone or in combination with LXXLL motif mutations (L/A). Scale bar: 10µm.

**Fig. S5.**
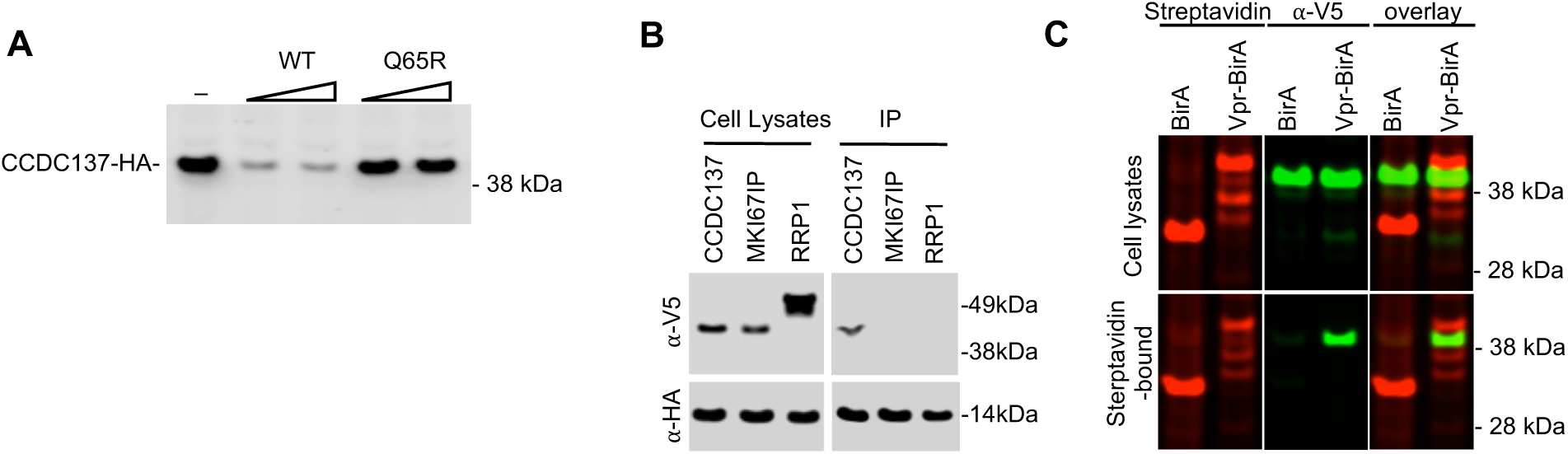
Physical interaction between Vpr and CCDC137. (A) 293T cells were transfected with HA-tagged CCDC137 expression plasmid along with varying amounts (0 ng, 25 ng, or 50 ng) of HA-tagged wild-type NL4-3 Vpr (WT), mutant Q65R (Q65R), expression plasmids. Twenty-eight hours post transfection, cells were harvested for Western blot analysis. (B) Western blot analysis of cell lysates and immunoprecipitates (IP) following cotransfection of 293T cells with plasmids expressing V5-tagged CCDC137, MKI67IP, or RRP1 along with a HA-Vpr expression plasmid. Immunoprecipitation was performed with an anti-HA monoclonal antibody and protein G beads. (C) Western analysis of cell lysates and streptavidin bead-bound proteins following transfection of 293T cells stably expressing V5 tagged CCDC137 with plasmids expressing BirA (R118G) or Vpr-BirA (R118G). Cells were treated with 50 µM biotin at 24 hours after transfection, and 10 µM MG132 at 40 hours post transfection, and harvested at 44 hours post transfection. Biotinylated proteins were detected with IRDye 680RD Streptavidin (red, left panel) and CCDC137 was detected with *α*-V5 antibody (green, middle panel).

**Fig. S6.**
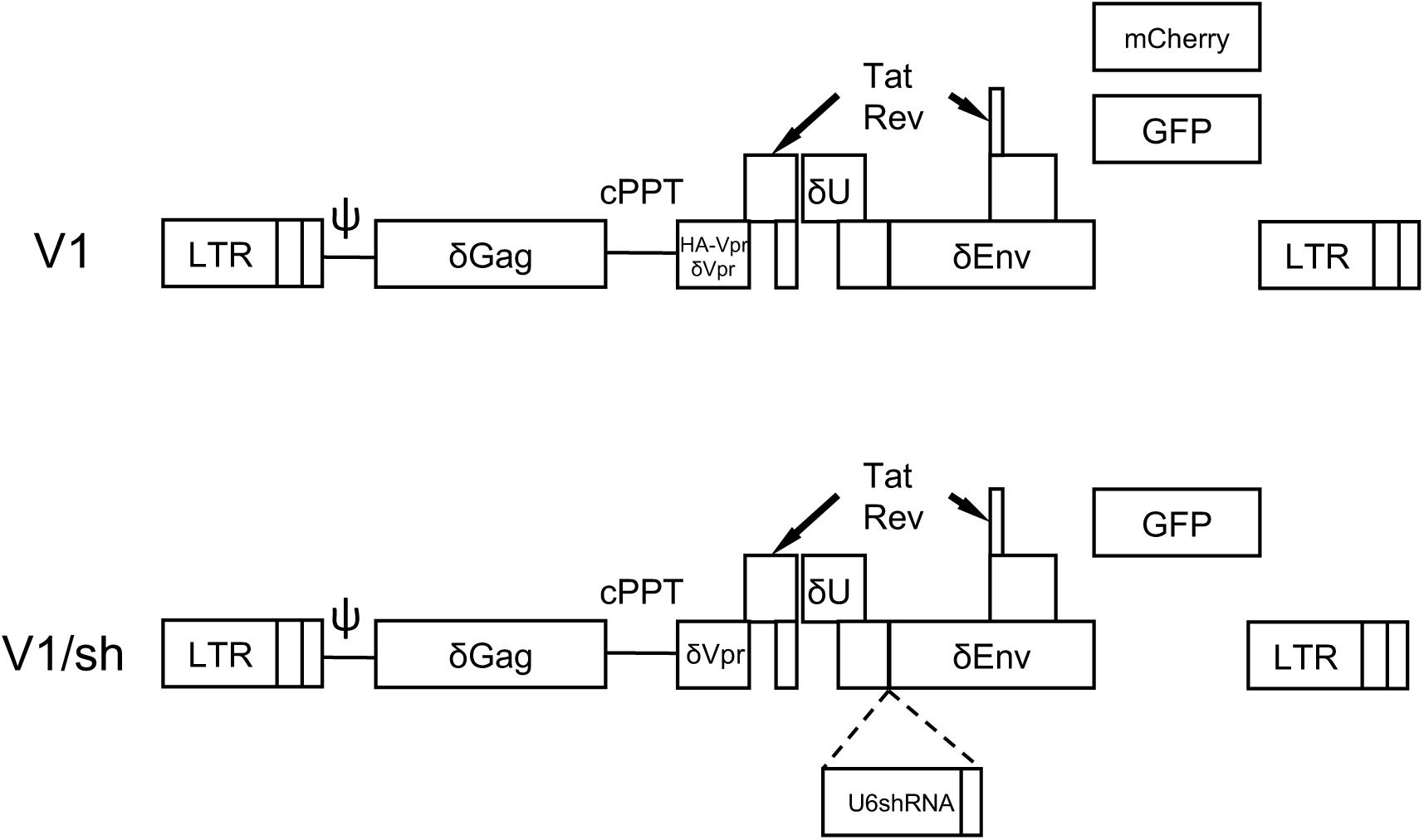
V1-based minmal HIV-1 genomes. Schematic representation of the minimal HIV-1 genomes used herein that contain GFP or mCherry reporter genes to mark infected cells, and assess HIV-1 gene expression using live cell imaging or FACS. Large deletions are introduced into the Gag, Pol, Vif and Env ORFs, while the Vpu ORF contains a premature termination codon. In V1/δVpr, the Vpr ORF contains a deletion while in V1/HA-Vpr an intact Vpr ORF from HIV-1_NL4-3_, is present. A variant of V1, termed V1/sh contains an expression cassette for an shRNA driven by a U6 promoter.

**Fig. S7.**
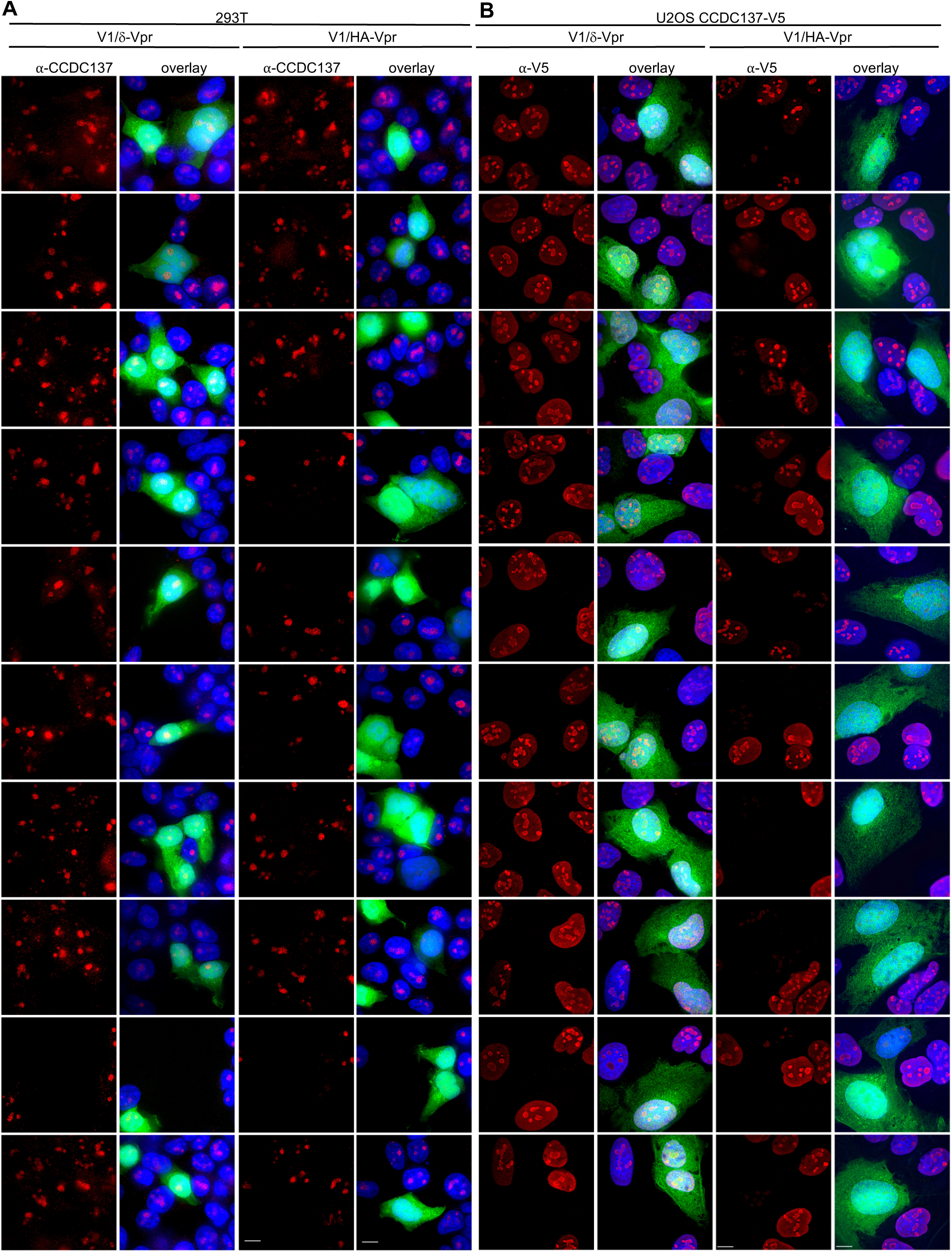
Immunofluorescent detection of CCDC137 depletion by Vpr. (A, B) Gallery of images following immunofluorescent staining to detect endogenous CCDC137 in 293T cells4 (A) or ectopic V5-tagged CCDC137 in U2OS cells (B) at 48 hours after infection with V1/δ-Vpr (*left*) or V1/HA-Vpr (*right*) at low MOI. Scale bar: 10 μm.

**Fig. S8.**
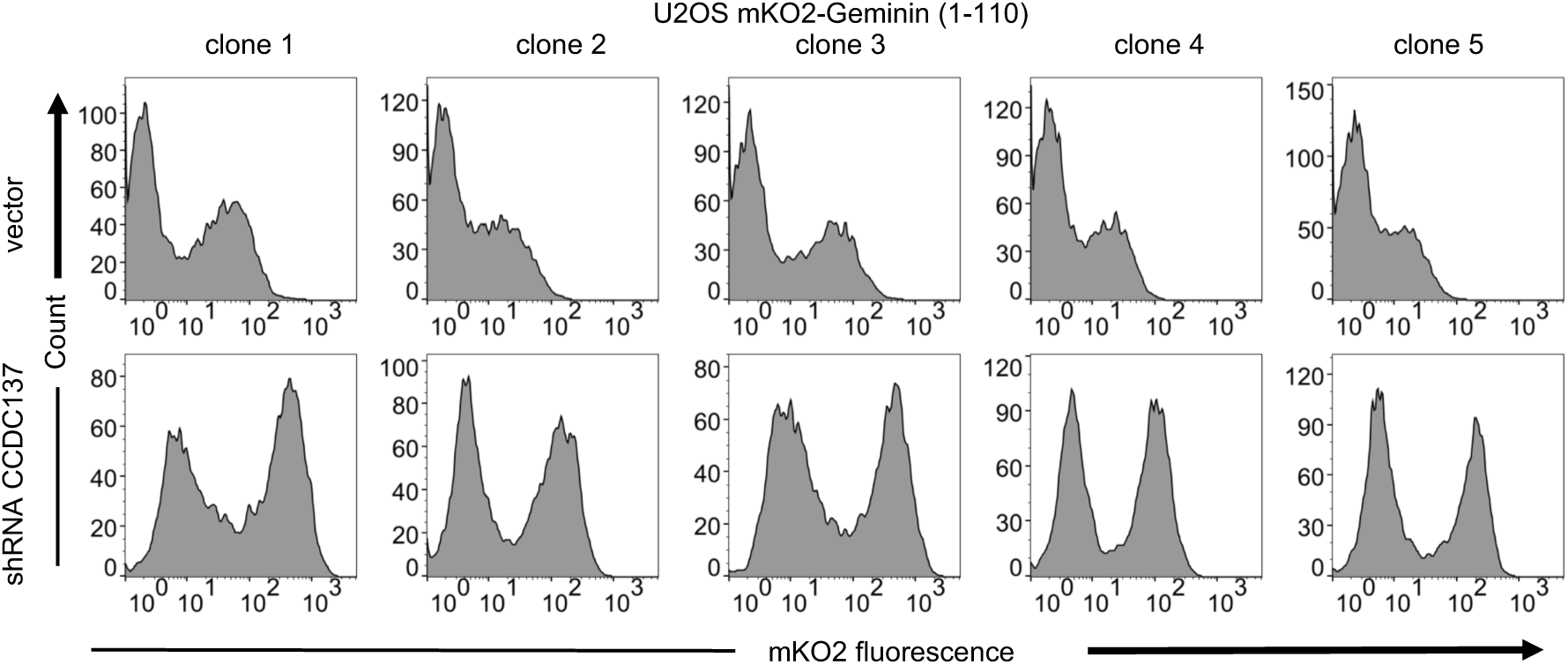
Cell cycle arrest induced by CCDC137 depletion. FACS analysis of U2OS-derived cell clones stably expressing an mKusabira-Orange2 (mKO2)-hGeminin (1-110 aa) fusion protein, following transduction with lentiviruses carrying no shRNA (vector, upper panels) or a CCDC137-targeting shRNA I (lower panels) and selection with puromycin for 40 hours.

**Fig. S9.**
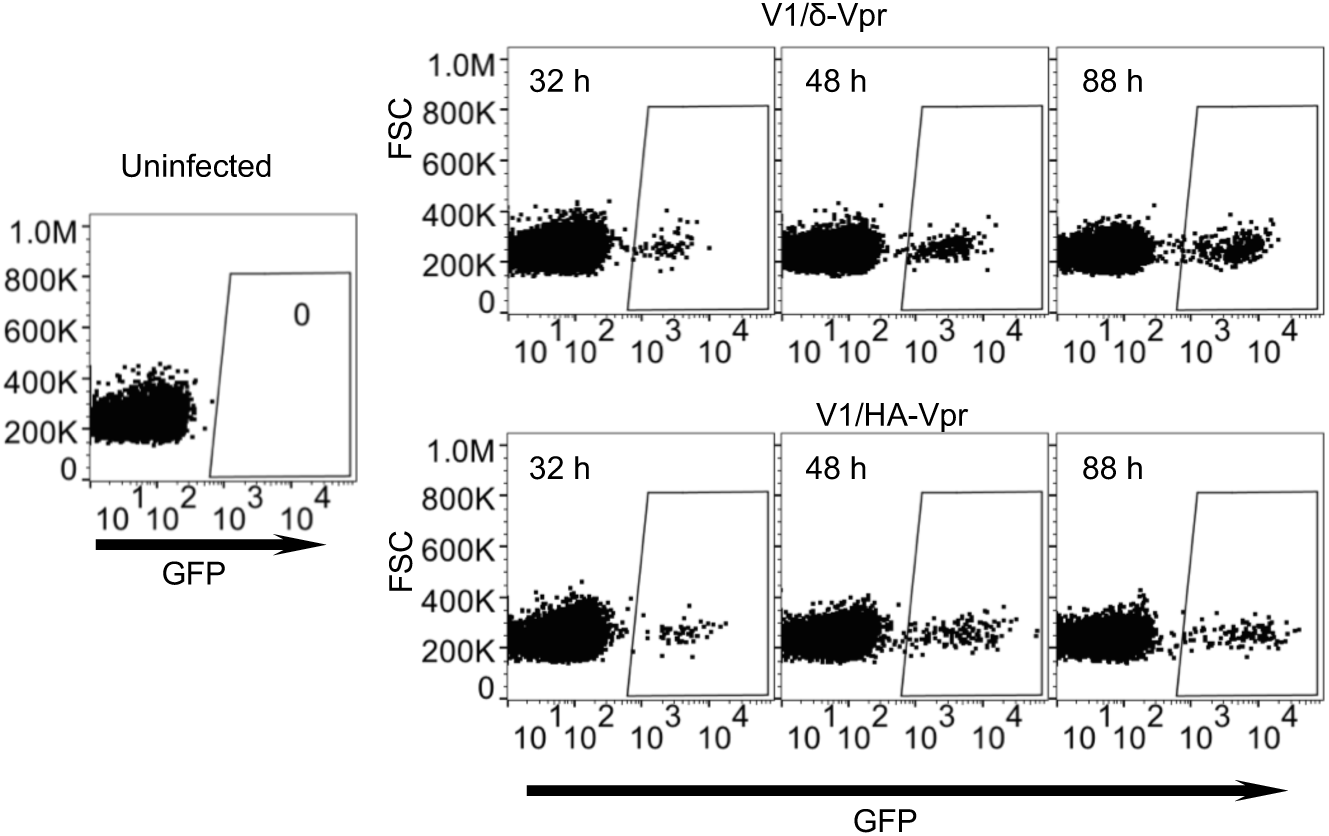
Enhancement of HIV-1 gene expression in CD4+ T-cells by Vpr. FACS analysis of GFP expression in activated primary CD4+ cells after infection with V1/δ-Vpr or V1/HA-Vpr. A representative donor is shown.

**Fig. S10.**
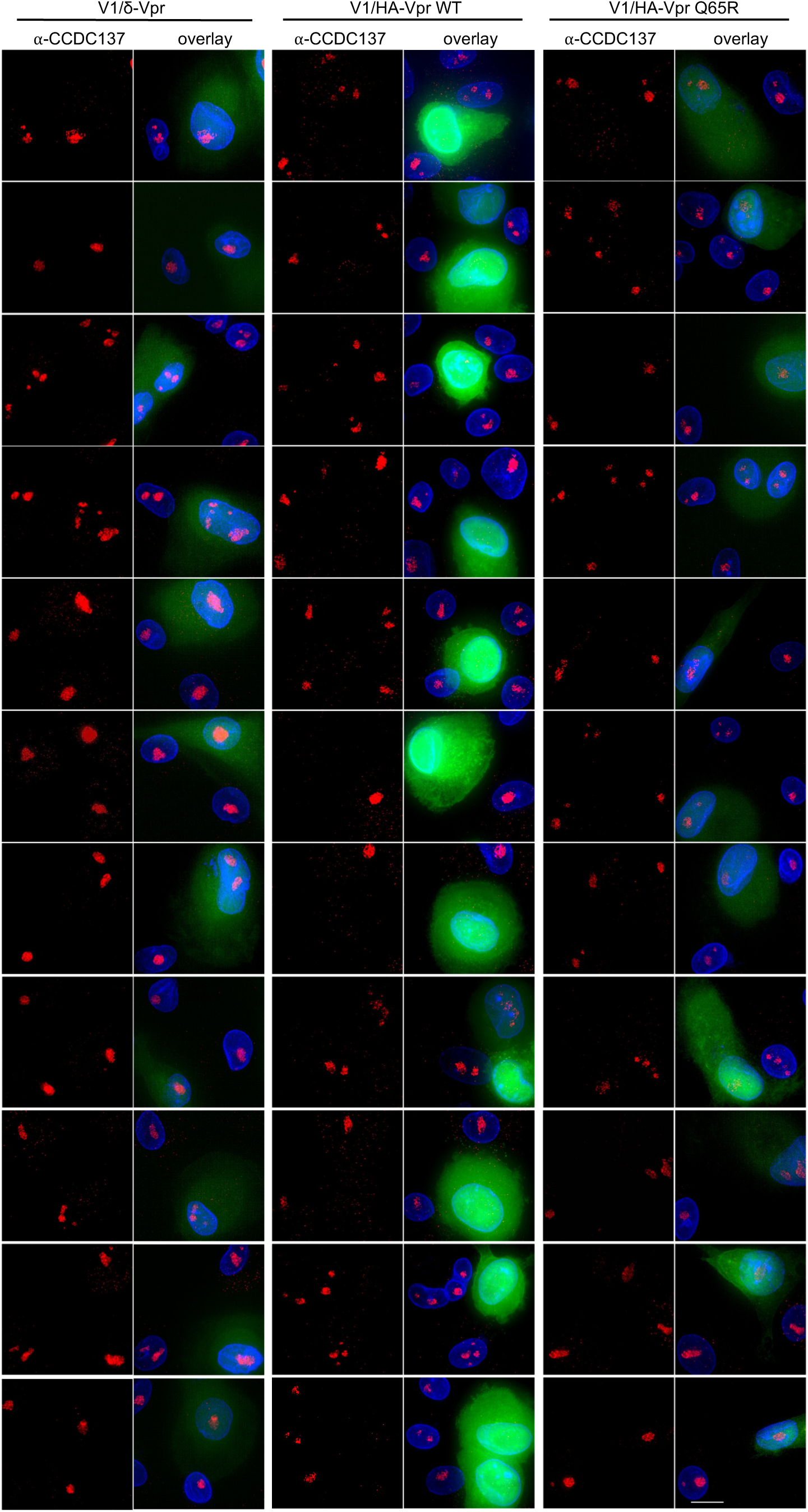
Immunofluorescent detection of CCDC137 depletion by Vpr and increased GFP expression in macrophages. (A, B) Gallery of images following immunofluorescent staining to detect endogenous CCDC137 and GFP expression in primary macrophages at 48 hours after infection with V1/δ-Vpr (*left*), V1/HA-Vpr (*center*) or V1/HA-Vpr (Q65R) (*right*) at low MOI. Scale bar: 10 μm.

**Fig. S11.**
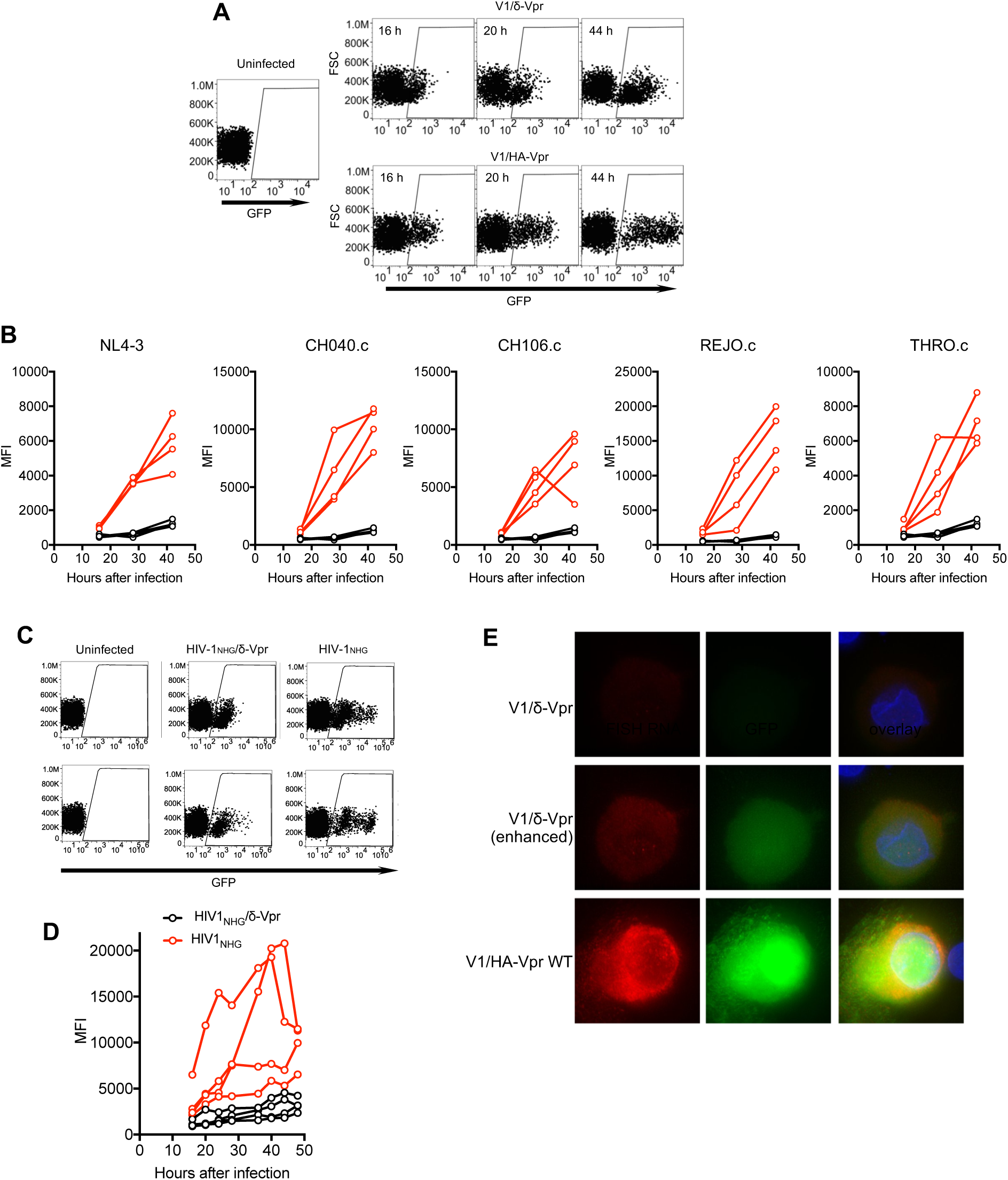
Enhancement of HIV-1 gene expression in macrophages by Vpr. (A) FACS analysis of GFP expression in primary macrophages after infection with V1/δ-Vpr or V1/HA-Vpr. A representative donor is shown (B) FACS analysis of GFP expression macrophages after infection with V1/δ-Vpr (Black) or V1/HA-Vpr (Red) derivatives encoding Vpr proteins from several different transmitted founder virus strains. The mean fluorescent intensity (MFI) of infected cells, gated as in (A) for 4 macrophage donors is plotted. (C) FACS analysis of GFP expression in primary macrophages after infection with HIV-1_NHG_/δ-Vpr or HIV-1_NHG_. A representative donor is shown. (D) FACS analysis of GFP expression in primary macrophages after infection with HIV-1_NHG_/δ-Vpr (Black) or HIV-1_NHG_ (Red). The mean fluorescent intensity (MFI) of infected cells, gated as in (A) for 4 macrophage donors is plotted. (E) Representative images of primary macrophages infected with V1/δ-Vpr or V1/HA-Vpr and subjected to fluorescent in situ hybridization (FISH) based detection of HIV-1 RNA using probes directed at the GFP sequence. The FISH signal is displayed in red and GFP protein signal is displayed in green. The upper (V1/δ-Vpr) and lower (V1/HA-Vpr) rows are displayed with the same gain and brightness/contrast settings. The center (V1/δ-Vpr) row is a duplicate of the upper row, displayed with enhanced brightness to enable visualization of the FISH and GFP signals.

**Fig. S12.**
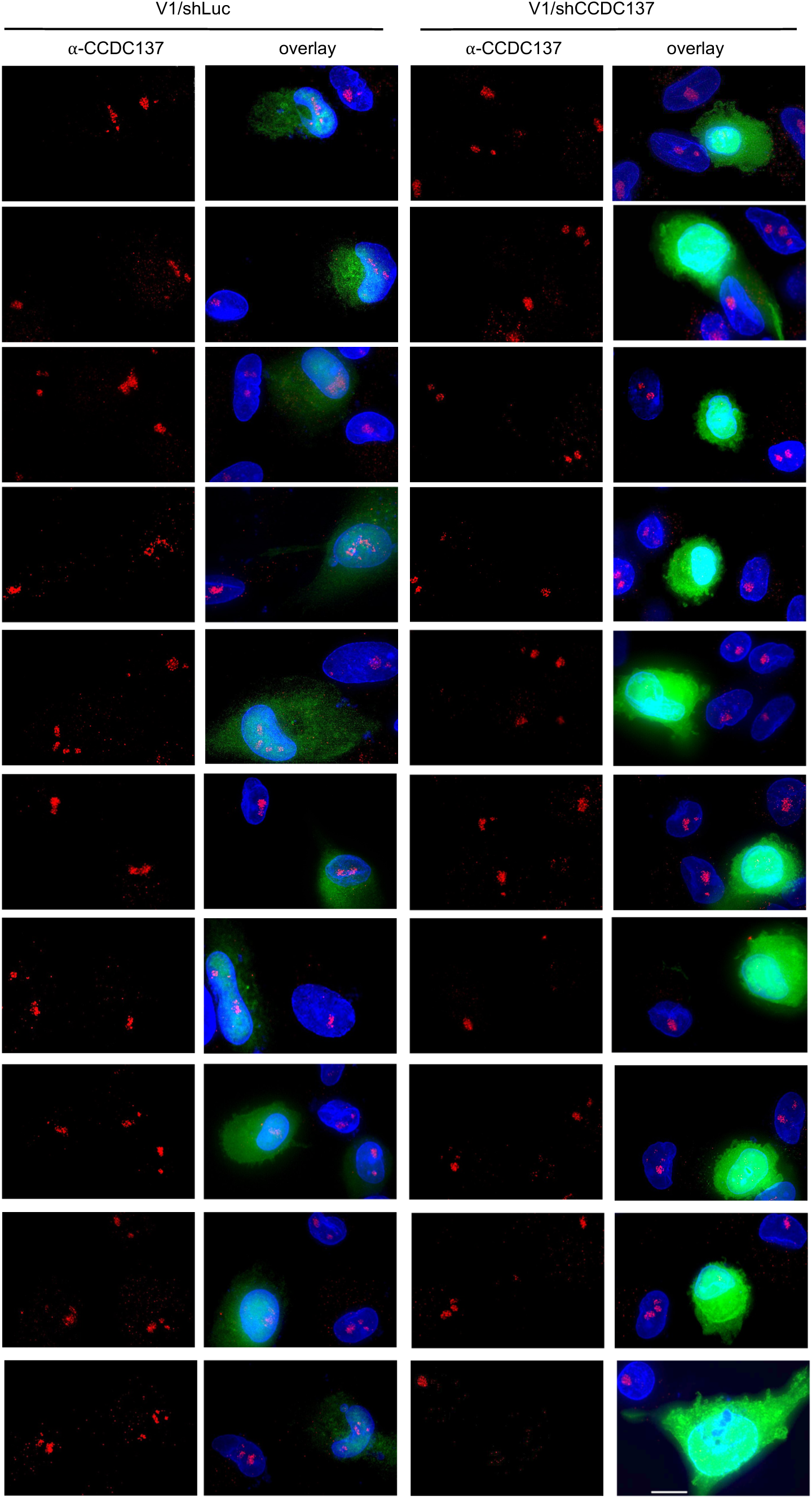
Immunofluorescent detection of CCDC137 depletion by shRNA and increased GFP expression in macrophages. Gallery of immunofluorescent staining to detect GFP expression as well as endogenous CCDC137 in primary macrophages at 48 hours after infection with V1/sh (*left*) or V1/shCCDC137 II (*right*) at low MOI. Scale bar: 10 μm.

**Fig. S13.**
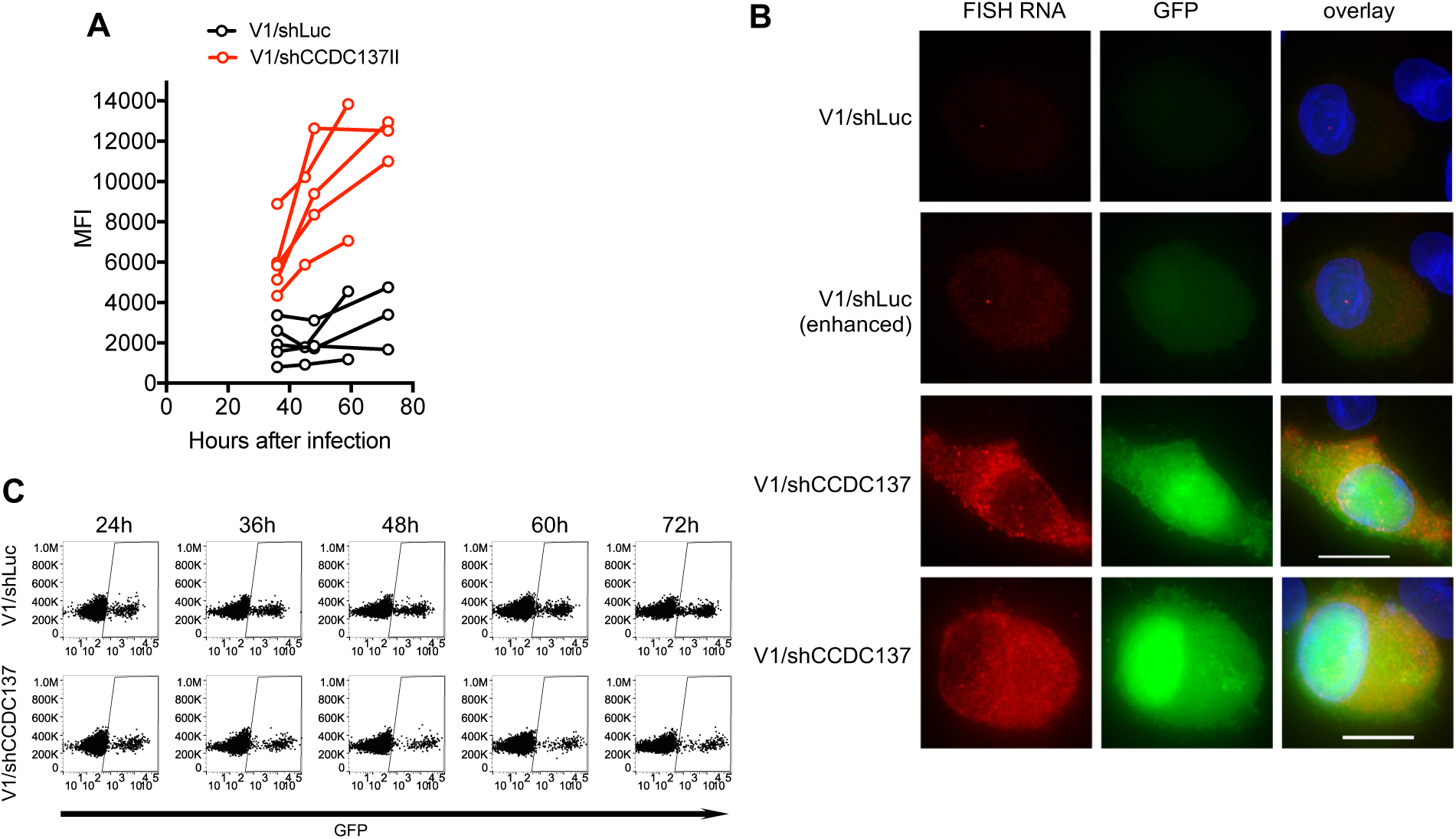
Enhancement of HIV-1 gene expression in macrophages and CD4+ T-cells by shRNA-mediated CCDC137 depletion. (A) FACS analysis of GFP expression in macrophages from four additional donors after infection with V1/shLuc or V1/shCCDC137. MFI of infected cells, gated as in Fig 4F is plotted. (B) Representative images of primary macrophages infected with V1/shLuc or V1/shCCDC137II and subjected to fluorescent in situ hybridization (FISH) based detection of HIV-1 RNA using probes directed at the GFP sequence. The FISH signal is displayed in red and GFP protein signal is displayed in green. The upper (V1/shLuc) and lowest two (V1/shCCDC137II) rows are displayed with the same gain and brightness/contrast settings. The second (V1/shLuc) row is a duplicate of the upper row, displayed with enhanced brightness to enable visualization of the FISH and GFP signals. (C) FACS analysis of GFP levels in primary CD4+ cells at the indicated time points after infection with V1/shLuc or V1/shCCDC137. A representative donor is shown.

**Movie S1. G2/M arrest induced by CCDC137.**

U2OS expressing mClover-hGeminin (1-110 aa) were transduced with a pLKO lentivirus vector containing no shRNA (upper row), or shRNA targeting CCDC137 (lower row). Images are phase contrast (left), mClover-hGeminin (green, center) and an overlay (right), and were acquired commencing at 36 hours post transduction, and every 30 minutes for the subsequent 72 hours.

**Movie S2. G2/M arrest and enhanced HIV-1 gene expresson induced by Vpr.**

U2OS cells expressing mClover-hGeminin (1-110 aa) were infected with V1/δ-Vpr (upper row) or V1/HA-Vpr (lower row) carrying mCherry. Images (from left to right) are phase contrast, mCherry (red), mClover-hGeminin (green) and an overlay of all channels, and were acquired commencing 18 hours after viral infection, and images every 30 minutes for the subsequent 48 hours.

**Movie S3. HIV-1 (V1/mCherry) expression in CCDC137 depleted U2OS cells.**

U2OS expressing mClover-hGeminin (1-110 aa) were transduced with lentiviruses containing no shRNA (upper row), or shRNAs targeting CCDC137 (lower row). At 28 hours post transduction, cells were infected with V1/δ-Vpr carrying mCherry. Images (from left to right) are phase contrast, mCherry (red), mClover-hGeminin (green) and an overlay of all channels, and were acquired commencing 12 hours after V1/δ-Vpr/mCherry infection and every 30 minutes for the subsequent 60 hours.

**Movie S4. HIV-1 (V1/δ-Vpr vs V1/HA-Vpr) expression in macrophages (donor #1).**

Freshly prepared human macrophages were infected with V1/δ-Vpr (upper row) or V1/HA-Vpr (lower row) carrying GFP. Images are phase contrast (left), GFP (green, center) and an overlay (right), and were acquired commencing at 24 hours after viral infection, and every 30 minutes for the subsequent 60 hours.

**Movie S5. HIV-1 (V1/δ-Vpr vs V1/HA-Vpr) expression in macrophages (donor #2).**

Freshly prepared human macrophages were infected with V1/δ-Vpr (upper row) or V1/HA-Vpr (lower row) carrying GFP. Images are phase contrast (left), GFP (green, center) and an overlay (right), and were acquired commencing at 24 hours after viral infection, and every 30 minutes for the subsequent 60 hours.

**Movie S6. HIV-1 (V1/shLuc vs V1/shCCDC137) expression in macrophages (donor #3).** Freshly prepared human macrophages were infected with V1/shLuc (upper row) or V1/shCCDC137 (lower row) carrying GFP. Images are phase contrast (left), GFP (green, center) and an overlay (right), and were acquired commencing at 20 hours after viral infection, and every 30 minutes for the subsequent 38 hours.

**Movie S7. HIV-1 (V1/shLuc vs V1/shCCDC137) expression in macrophages (donor #4).** Freshly prepared human macrophages were infected with V1/shLuc (upper row) or V1/shCCDC137 (lower row) carrying GFP. Images are phase contrast (left), GFP (green, center) and an overlay (right), and were acquired commencing at 20 hours after viral infection, and every 30 minutes for the subsequent 38 hours.

**Data S1.**

List of proximal/interacting proteins identified using Mass Spectrometry analysis of biotinylated proteins in BirA (R118G)-fused Vpr-expressing cells. The dataset includes UniProtKB accession number, description, mean peak area, scores (the sum of the highest ions score for each distinct peptide), percent coverage (calculated by dividing the number of amino acids in all found peptides by the total number of amino acids in the entire protein sequence), the number of distinct peptides used for identification in a protein, and peptide spectrum matches (PSM, the total number of identified peptide sequences for the protein). The dataset contains results from duplicate samples.

